# Apurinic/apyrimidinic nuclease 1 drives genomic evolution contributing to chemoresistance and tumorigenesis in solid tumor

**DOI:** 10.1101/2022.04.20.488830

**Authors:** Subodh Kumar, Jiangning Zhao, Srikanth Talluri, Leutz Buon, Shidai Mu, Bhavani Potluri, Chengcheng Liao, Jialan Shi, Chandraditya Chakraborty, Gabriel B. Gonzalez, Yu-Tzu Tai, Jaymin Patel, Jagannath Pal, Hiroshi Mashimo, Mehmet K. Samur, Nikhil C. Munshi, Masood A. Shammas

**Affiliations:** Dana Farber Cancer Institute, Boston, MA 02215; Veterans Administration Boston Healthcare System, West Roxbury, MA 02132; Harvard Medical School, Boston, MA 02215; Beth Israel Deaconess Medical Center, Boston, MA 02215; Pt. J.N.M. Medical College, Raipur, Chhattisgarh, India

**Keywords:** Genomics, Genomic Instability, Genomic Evolution, Homologous Recombination, Esophageal Adenocarcinoma, Solid Tumor, Tumorigenesis, APE1, APEX1

## Abstract

Genomic instability fuels genomic alterations that befit cancer cells with necessary adaptations to keep proliferating and overcome the impact of host anti-tumor immunity and cytotoxic therapy. Since DNA breaks are required for genomic rearrangements to take place, we hypothesized that dysregulated nuclease activity mediates genomic instability in cancer. Using an integrated genomics protocol, we identified a four gene deoxyribonuclease signature correlating with genomic instability in six human cancers which included adenocarcinomas of esophagus (EAC), lung, prostate, stomach, pancreas and triple negative breast cancer. Functional screens confirmed the role of these nucleases in genomic instability and growth of cancer cells. Apurinic/apyrimidinic nuclease 1 (APE1), identified as top nuclease in functional screen, was further investigated in five cell lines representing four solid tumors (EAC, lung, prostate and breast cancer). We demonstrate that chemical as well as transgenic suppression of *APE1* impaired growth/colony formation and increased cytotoxicity of chemotherapeutic agent, whereas inhibited spontaneous as well as chemotherapy-induced DNA breaks, homologous recombination (HR) activity and genomic instability in all cancer cell types tested. Treatment with APE1 inhibitor also impaired tumor growth and significantly increased efficacy of a chemotherapeutic agent in a subcutaneous mouse model of EAC. Overexpression of APE1 in normal esophageal epithelial cells increased DNA breaks and HR activity, leading to massive mutational, copy number as well as karyotypic instability. Evaluation of by whole genome sequencing identified HR as the top mutational process activated by APE1. Normal cells overexpressing APE1 grew as tumors in mice and tumors removed from mice displayed additional karyotypic changes, providing evidence of genomic instability *in vivo*. Overall, our data demonstrate that elevated APE1 dysregulates HR activity, G2/M checkpoint and genome stability thus contributing to tumorigenesis and chemoresistance in cancer. Therefore, inhibitors of *APE1* have potential to inhibit growth and increase cytotoxicity of chemotherapeutic agents while minimizing spontaneous as well as chemotherapy-induced genomic damage and instability in EAC and other solid tumors.

## INTRODUCTION

Most cancers display diverse genomic aberrations^1–6^ which evolve over time^3,7–9^, indicating a marked instability at DNA and chromosomal levels^1–13^. Genomic instability is a relatively early event during oncogenesis and has been observed even in pre-neoplastic lesions^3,7,8^. Ongoing changes at DNA and chromosome levels provide new characteristics for growth and survival as well as enable affected cells to overcome immune surveillance^14^ and contribute to disease progression^11,13,15^. Genomic instability underlies clonal evolution and tumor heterogeneity, and increased tumor heterogeneity can in turn lead to chemoresistance and relapse^16–17^.

Esophageal adenocarcinoma (EAC), a cancer associated with gastroesophageal reflux, arises from Barrett’s esophagus (BE), a precancerous condition which progresses to cancer through advancing stages of dysplasia^18^. EAC genome is strikingly aberrant^19^. Genomic instability, which exists in EAC at precancerous stage^3,8,18–24^, seems to increase over time^3,23^ and contribute to development of cancer and its progression. Consistently, the cancer is mostly chemoresistant^25^ and associated with poor prognosis^26^.

Rate of mutation in lung cancer has been estimated to be the second highest among cancers, indicating a striking genomic instability which gives rise to a heterogenous genetic landscape^27^. A signature, comprised of seven long non-coding RNAs, which correlated with genomic instability, was able to prognosticate overall survival of lung adenocarcinoma patients^28^. A striking genomic instability, as evident from complex genomic aberrations including chromothripsis, detected in prostate cancer patients has been linked to cancer progression^29^. Genomic instability has also been implicated in etiology and progression of breast cancer^30^. Defects in the pathway intermediates ensuring DNA repair and proper segregation of chromosomes seem to be among prominent mechanisms contributing to genomic instability in breast cancer^31^. Luminal B tumors of breast, that are positive for estrogen receptor but negative for progesterone receptor, have increased genomic instability and demonstrate increased growth rate and tendency to develop resistance to tamoxifen^32^. Similarly, it has been demonstrated that a genomic instability-related score calculated based on copy number alterations can predict prognosis in luminal A breast cancer^33^. It is now becoming quite evident that genomic instability is a promising target in cancer^34,35^. Chromosomal instability has been implicated in cancer progression and development of resistance to treatment and seems to associate with poor clinical outcome^13^. Identification of genes and pathways which drive genomic instability and evolution will not only improve our understanding of the oncogenic process but will help develop better strategies to treat and/or prevent cancer.

Cells in our body are constantly exposed to a number of DNA-damaging agents which could be of exogenous or endogenous origin. This leads to a large number and types of DNA lesions which occur in our cells on daily basis^36,37^. These include DNA strand breaks, cross-linking and missing altered or incorrect bases. In normal cellular condition, all these lesions are accurately repaired on daily basis. If the lesions, especially the double-strand breaks, are not repaired in G2 and persist into mitosis they can potentially lead to chromosomal aberrations or rearrangements. In normal cells, multiple DNA repair and related pathways coordinate to efficiently and accurately repair DNA lesions. Homologous recombination (HR), a repair pathway dependent on extensive sequence similarity between damaged and template DNA strands, is known to be the most precise DNA repair system^38,39^. The HR pathway contributes to the repair of several types of DNA lesions which include double-stranded breaks, single-stranded gaps and strand crosslinks thus ensuring the maintenance of genomic integrity and stability^40^. Accurate repair of DNA lesions and the maintenance of genome stability and integrity is strongly dependent on cell cycle checkpoints. In this regard, G2 along with the later part of S and earlier part of M phases, has a special significance. This is because in this phase of cell cycle, the HR pathway can utilize the sister chromatid as template to ensure error-free or most accurate repair of DNA lesions^41–43^. However, precise and error-free nature of HR depends on its strict regulation. Dysregulation of HR, whether its reduced or increased activity, poses risk to genome stability^42^. Our data in EAC and multiple myeloma demonstrate that spontaneously elevated HR activity contributes to genetic instability in these cells^11–12^. Dysregulated HR is also attributed to drug resistance^11^ and growth of cancer cells in subcutaneous tumor model^44^.

Purpose of this study was to identify genes driving genomic instability and evolution in EAC and possibly other solid tumors. Since DNA breaks are required for rearrangements to take place, we hypothesized that elevated deoxyribonuclease activity drives genomic instability in cancer. To identify such nucleases, we first identified those whose expression correlated with genomic instability in six different solid tumors including EAC. Expression of four nucleases correlated with genomic instability in all six cancers. Evaluation of these four nucleases in the loss and gain of function screens identified apurinic/apyrimidinic nuclease 1 (APE1), a major base excision repair protein, to have the strongest overall impact on genomic instability and growth of cancer cells. APE1 (apurinic/apyrimidinic endonuclease 1) plays a key role in the initiation of base excision repair by cleaving the DNA at 5’ of abasic site^45^. Loss of a base leading to emergence of an abasic site in DNA can occur either spontaneously or by activity of DNA glycosylases which have ability to remove damaged bases from DNA. Loss of a base occurs frequently in a cell^46–48^. However, thousands of such lesions generated on a daily basis are fixed because of an efficient repair system^49–50^. Our data in multiple myeloma has demonstrated that APE1 also regulates HR activity through its role in transcriptional regulation of recombinase RAD51^51^. In this study, we investigated the role of APE1 in genomic instability and growth in five cell lines representing four solid tumors (EAC, lung, prostate and breast cancer). Using cancer and normal cell types as well as in vitro and in vivo model systems, we demonstrate that elevated APE1 activity drives genomic instability contributing to tumorigenesis and chemoresistance. The ability of APE1 to dysregulate DNA repair and G2/M checkpoint could be attributed to these processes. Inhibitors of *APE1* have potential to inhibit growth and increase cytotoxicity of chemotherapeutic agents while minimizing spontaneous as well as chemotherapy-induced genomic damage and instability in EAC and other solid tumors.

## MATERIALS AND METHODS

### Patient datasets and specimens

The Cancer Genome Atlas (TCGA) data were used to investigate gene expression and copy number events in six human cancers. These included esophageal adenocarcinoma (EAC), stomach adenocarcinoma, prostate adenocarcinoma, pancreatic adenocarcinoma, triple negative breast cancer, and lung adenocarcinoma. De-identified specimens of normal esophageal epithelial squamous (NES), Barrett’s esophagus (BE, a precancerous lesion for EAC), dysplasia and EAC were provided by our collaborator Dr. Hiroshi Mashimo, who has an active protocol (R&D #2490, ID #1578027) at Boston VA Healthcare Center, West Roxbury, MA. Esophageal cancer progression tissue arrays were purchased from US Biomax, Inc. (Rockville, MD).

### Antibodies

Affinity purified human APE1 antibody (NB100-101), Novus Biologicals LLC, Centennial, CO; FEN1 Antibody (2746s), Cell Signaling Technology, Inc., Danvers, MA; EXO1 antibody (A302-639A), Bethyl Laboratories, Montgomery, TX, USA; EME1 antibody (PA5-101988), Invitrogen, Carlsbad, CA, USA; RAD51 antibody (ab176458), Abcam, Cambridge, MA, USA; RPA32(A300-244A), Bethyl Laboratories, Montgomery, TX, USA; RPA32 (phospho Ser4 and phospho Ser8) antibody, Novus Biologicals LLC, Centennial, CO; Anti-phospho-H2A.X (Ser139) antibody, (07-164), Sigma-Aldrich, St. Louis, MO, USA; GAPDH (14C10) rabbit monoclonal antibody ( #2118), Cell Signaling Technology, Inc., Danvers, MA; β-Tubulin antibody (#2146), Cell Signaling Technology, Inc., Danvers, MA.

### Overexpression and knockdown plasmids and siRNAs

Lentivirus-based shRNAs targeting APE1, lentivirus-based expression plasmids carrying open reading frames of FEN1 and APE1 and MISSION esiRNAs targeting human APEX1, EXO1, EME1 and FEN1 were purchased from Sigma-Aldrich, St. Louis, MO, USA. Lentivirus-based expression plasmids carrying open reading frames of EME1 and EXO1 were purchased from Abcam, Cambridge, MA, USA.

### Identification of a deoxyribonuclease signature correlating with genomic instability

We hypothesized that if elevated expression of a gene correlates with increased genomic instability in multiple human cancers, it could be a potential driver of genomic evolution. Since dysregulated nuclease activity can directly impact genome stability, we focused on deoxyribonucleases (excluding topoisomerases) for this study. To identify the nucleases whose expression correlated with genomic instability, we used following stepwise process: **1)** Investigated gene expression in normal and tumor samples for six human cancers in TCGA dataset and identified deoxyribonucleases (excluding topoisomerases); **2)** Assessed genomic instability in patient samples by counting total number of copy events in each patient; **3)** Integrated genomic instability data with expression data to identify deoxyribonucleases whose expression correlated with genomic instability; 4) Four gene nuclease signature was validated in functional screens. Apurinic/apyrimidinic nuclease 1 (APE1), demonstrating the strongest overall impact on genomic instability and growth, was evaluated in detail for its role in genomic instability, cell cycle and oncogenic process.

### Cell types

Normal human mammary epithelial cells were purchased from Lonza Bioscience (Walkersville, MD), normal primary human esophageal epithelial (HEsEpiC) cells were obtained from ScienCell Research Laboratories (Carlsbad, CA) and normal human diploid fibroblasts and human cancer cell lines — breast cancer (MCF7, MDA453), colon cancer (HT29, HCT116), epithelial lung carcinoma (A549) and prostate adenocarcinoma (PC3) purchased from American Type Culture Collection (Manassas, VA). Esophageal adenocarcinoma (EAC) cell lines (FLO-1, OE19 and OE33) were purchased from Sigma Aldrich Corporation (Saint Louis, MO). All cell types were cultured as reported previously^12,44, 52–54^.

### Cell Viability

Cell Titer-Glo Luminescent Viability Assay kit (Promega Corporation, Madison, WI) was used to assess cell viability.

### Modulation of gene expression/function

APE1 activity was modulated using chemical as well as transgenic manipulations. For chemical inhibition, APE1 Inhibitor III (API3), a specific inhibitor of APE1 (Axon Medchem LLC, Reston, VA) was used. For knockdown of APE1, EXO1, FEN1 and EME1 either esiRNAs or lentivirus-based shRNAs (Millipore/Sigma, Saint Louis, MO) were used. For overexpression, the lentivirus-based plasmid carrying APE1 gene under CMV promoter was used.

### Gene Expression Analysis and Biostatistics

Total RNA was isolated utilizing an “RNeasy” kit (Qiagen Inc., USA) and gene expression profile was evaluated using Human Gene 1.0 ST Array (Affymetrix, Santa Clara, CA) as described by us previously^11,12,52^.

### Homologous Recombination Assays

Homologous recombination (HR) activity was assessed using the plasmid-based functional assay described previously^12,44^. We also used a fluorescence-based homologous strand exchange (SE) assay to measure HR, as reported by Huang et al^55^.

### Detection of DNA breaks and end resection

DNA breaks were assessed by evaluating cells for γ-H2AX expression (a marker of DNA breaks), using Western blotting or by Comet assay, a gel-based method to visualize DNA breaks. Comet assay was done using OxiSelect™ Comet Assay Kit (Cell Biolabs Inc., San Diego, CA) as described by us previously^52^. DNA end resection, which is one of the important initial steps in HR, was assessed by investigating the phosphorylation of RPA32 on ser4 and ser^56–57^.

### Investigation of protein-protein interactions, expression and phosphorylation levels

EAC (FLO-1) cells were lysed and immunoprecipitated using anti-APE1 or anti-IgG antibodies and proteins in each complex identified using mass spectrometry. Proteins interacting with APE1 but not with IgG were identified and subjected to protein-protein interaction (PPI) network analysis using STRING (STRING; http://string-db.org; version 11.0). Interactions with a comprehensive enrichment score > 0.4 was considered to be statistically significant and visualized with Cytoscape 3.7.2 software. The network was clustered based on topology with Cytoscape’s plug-in MCODE (using parameters: MCODE score > 5, degree cutoff = 2, node score cutoff = 0.2, max Depth = 100, k-score = 2) to find densely connected regions and the most important modules in the PPI network. The cytoHubba plug-in was used to perform operations, and the Degree method was used to evaluate the key node genes. For investigation of protein expression and phosphorylation levels, the Cell Cycle Control/DNA Damage Phospho Antibody Arrays (Full Moon BioSystems, Inc., Sunnyvale, CA) containing highly specific and well-characterized antibodies were utilized. The antibodies on the array are site-specific and allow investigation of specific phosphorylation sites.

### Cell cycle analysis

Cells were washed twice with PBS and fixed in 95% ethanol. Cells were pelleted, washed and resuspended in propidium iodide/RNAse solution (Life Technologies) and investigated using flow cytometry. Data was analyzed using FLOJO software.

### Investigating genomic instability and evolution

Genomic instability and evolution were investigated using three different methods as described below:

#### Micronucleus assay

Micronuclei, one of the marker of genomic instability^58^, were investigated by a flow cytometry-based assay using a kit as previously reported by us^51,54^.

#### Single nucleotide polymorphism (SNP) arrays and Whole genome sequencing (WGS)

Briefly, the APEX1 was either suppressed (by shRNAs or small molecule) or transgenically-overexpressed, the cells were cultured for different durations, genomic DNA purified and analyzed using either SNP6.0 arrays (Affymetrix) or WGS. For all these experiments, the genome of “day 0” cells (harvested and saved in the beginning of each experiment) were used as baseline to identify changes in control and transgenically-modified/treated cells during their growth in culture. WGS and SNP data were analyzed as reported by us previously^11–12,54^.

### Investigating chromosomal instability

Normal and APE1-transformed human esophageal epithelial cells were prepared for cytogenetic analysis by treatment with colcemid (Gibco/Life Technologies, Grand Island, NY) at a final concentration of 0.1 ug/ml for 3 hr to accumulate cells in metaphase. Cells were then exposed to hypotonic (0.075 M KCl) solution for 25 minutes at 37°C and fixed with 3:1 methanol:acetic acid. Air-dried slides were made for mitotic index (MI) determination and FISH. MI was determined by counting a minimum of 500 cells from random microscope fields and by dividing the number of interphase (non-mitotic) cells by the number of metaphases (mitotic cells) in the same fields. MFISH was performed both with and without simultaneous myc FISH. SpectrumGreen-labeled myc probe (Vysis LSI MYC, Abbott Laboratories, Des Plaines, IL) was diluted 1:20 in ready-to-use MFISH probe (MetaSystems GmbH, Altlussheim, Germany). Hybridization and wash steps followed manufacturer’s instructions for MFISH with minor modifications. Slides were mounted in antifade solution with DAPI counterstain (Vectashield, Vector Labs). Hybridized slides were examined on an Olympus AX-70 microscope equipped with appropriate filters, and images captured with Isis (MetaSystems) imaging software.

### Evaluation of impact of APE1 overexpression on tumorigenesis in SCID mice

Normal human esophageal epithelial (HEsEpiC) cells were transfected with control plasmid (C) or the plasmids carrying APE1 gene under CMV promoter (APE1O) to induce APEX1 expression. Five-week-old male CB-17 SCID mice were purchased from Charles River Laboratories (Wilmington, MA) and maintained following guidelines of the Institutional Animal Care and Use Committee (IACUC); all experimental procedures were approved by the IACUC and the Occupational Health and Safety Department of Dana Farber Cancer Institute, Boston, MA. Cells carrying control plasmid (C) were injected on left side and APE1-overexpressing (APE1O) cells injected on right side of each mouse. Tumor sizes were measured twice a week and animals euthanized when tumors reached 2 cm^3^ in volume or when paralysis or major compromise in their quality of life occurred. Mice were then euthanized, tumors removed and evaluated for karyotypic changes. Tissue was minced and dissociated with vigorous pipetting (no collagenase was used) and set up for tissue culture in multiple flasks. Each flask was harvested for metaphase preparations when appropriate, according to observed cell growth with varying colcemid concentration and exposure time. Evaluable metaphases were obtained at day 4 and were used for this analysis.

## Acknowledgements/Funding

This work was supported by Department of Veterans Affairs Merit Review Award I01BX001584-01 (NCM), NIH grants P01-155258 and 5P50 CA100707 (MAS, MKS, NCM), Leukemia and Lymphoma Society translational research grant (NCM) and Guangxi Natural Science Foundation Program Grant 2018GXNSFBA281026 for postdoctoral training (CL).

## Author contributions

MAS envisioned the study, analyzed and interpreted data and prepared manuscript; NCM assisted in data interpretation and critical review of manuscript; LB and MKS conducted bioinformatic and statistical analyses; SK, JZ and ST equally contributed to major experiments and manuscript preparation; SM, BP, CL, JS, CC, GBG, YT and JP contributed to specific experiments and data analyses; HM provided specimens and assisted in interpretation and critical review of manuscript.

## RESULTS

### Identification and functional validation of deoxyribonuclease signature correlating with genomic instability in solid tumors

Genomic instability is a prominent feature of cancer cells. Since DNA must be cut and/or processed for genomic rearrangements to take place, we hypothesized that dysregulated nuclease activity mediates genomic instability in cancer. Consistent with this hypothesis, the evaluation of **γ**H2AX expression in nine cancer cell lines representing five human cancers (esophageal adenocarcinoma, breast cancer, colon cancer, epithelial lung carcinoma and prostate adenocarcinoma) indicated that spontaneous DNA breaks are increased in cancer relative to normal cell types (human mammary epithelial cells, human diploid fibroblasts, human esophageal epithelial cells) (**Supplementary Figure 1**). To identify the nucleases contributing to spontaneous DNA breaks and genomic instability, we used an integrated genomics protocol followed by functional validation (Figure 1a). **<u>Identification</u>:** Gene expression and copy number data of six human cancers (esophageal adenocarcinoma, EAC; lung adenocarcinoma, LUAD; prostate adenocarcinoma, PRAD; stomach adenocarcinoma, STAD; pancreatic adenocarcinoma, PAAD; and triple negative breast cancer, TNBC) from TCGA dataset were used in a stepwise process as shown in Figure 1a (panel I). Genomic instability in each patient sample in each cancer was assessed by counting total copy number events per patient; data from triple negative breast cancer (TNBC) patients is shown as an example in Figure 1a (panel I). Copy number events in patient samples of other cancer types are shown in **Supplementary Figure 2.** Genomic instability data were then integrated with expression data to identify deoxyribonucleases whose expression correlated with genomic instability. Four deoxyribonucleases (FEN1, EXO1, EME1 and APE1) correlated with genomic instability in all six cancers (Figure 1a, panels II-III). We thus identified four gene deoxyribonuclease genomic instability signature (N-GIS) representing six human cancers. **<u>Functional validation</u>:** For functional validation, deoxyribonuclease genomic instability signature (N-GIS) genes (APE1, EXO1, FEN1 and EME1) were suppressed in EAC (FLO1) cells using esiRNAs, knockdown confirmed by Western blotting (Figure 1b, panel I) and impact on genome stability and cell viability assessed. Genomic instability was assessed by evaluation of micronuclei (a marker of genomic instability)^58^. Representative images of nuclei and micronuclei (Figure 1b, panel II) and bar graph summarizing the results from three independent experiments (Figure 1b, panel III) are shown.

**Figure 1.**
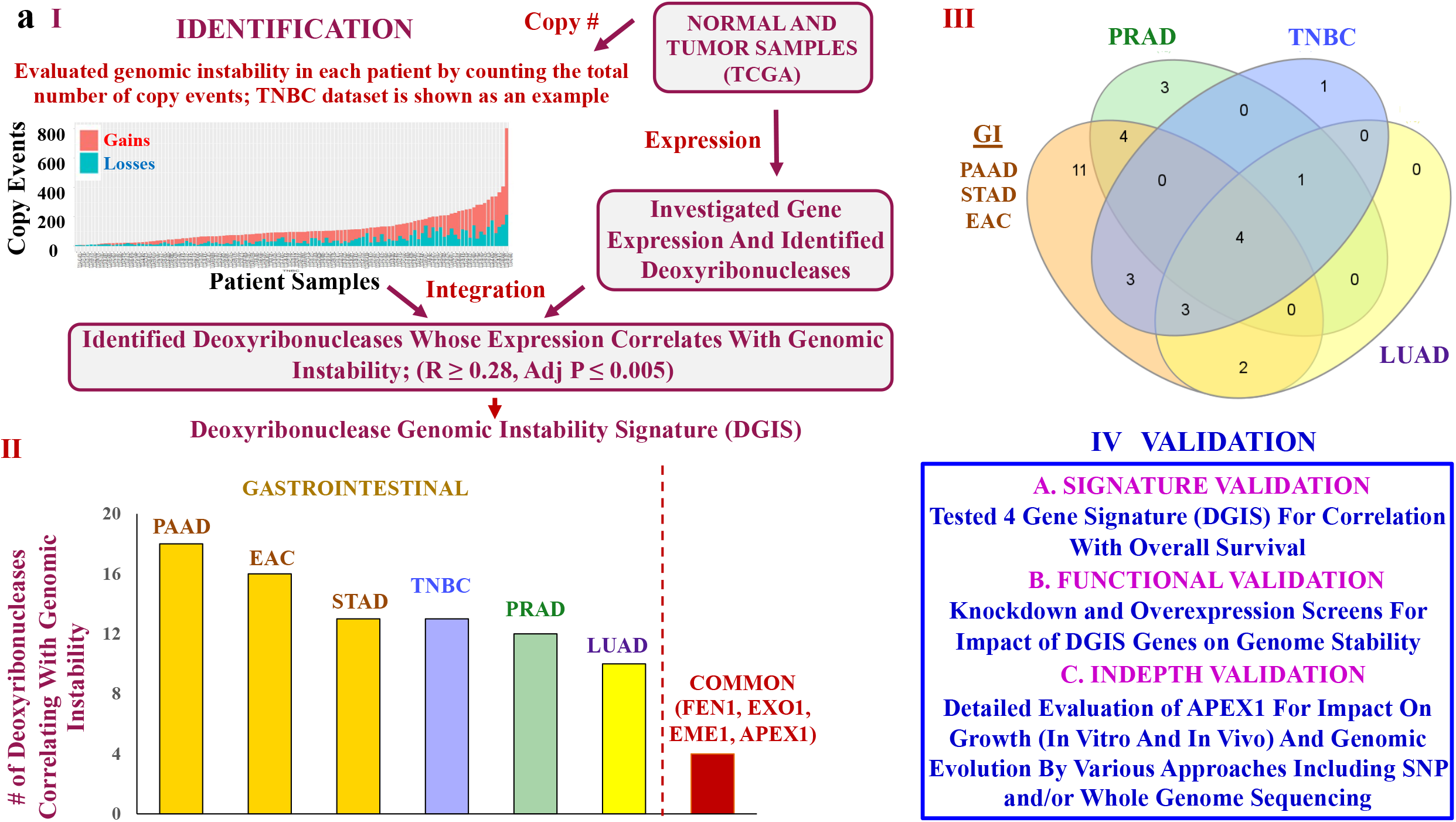
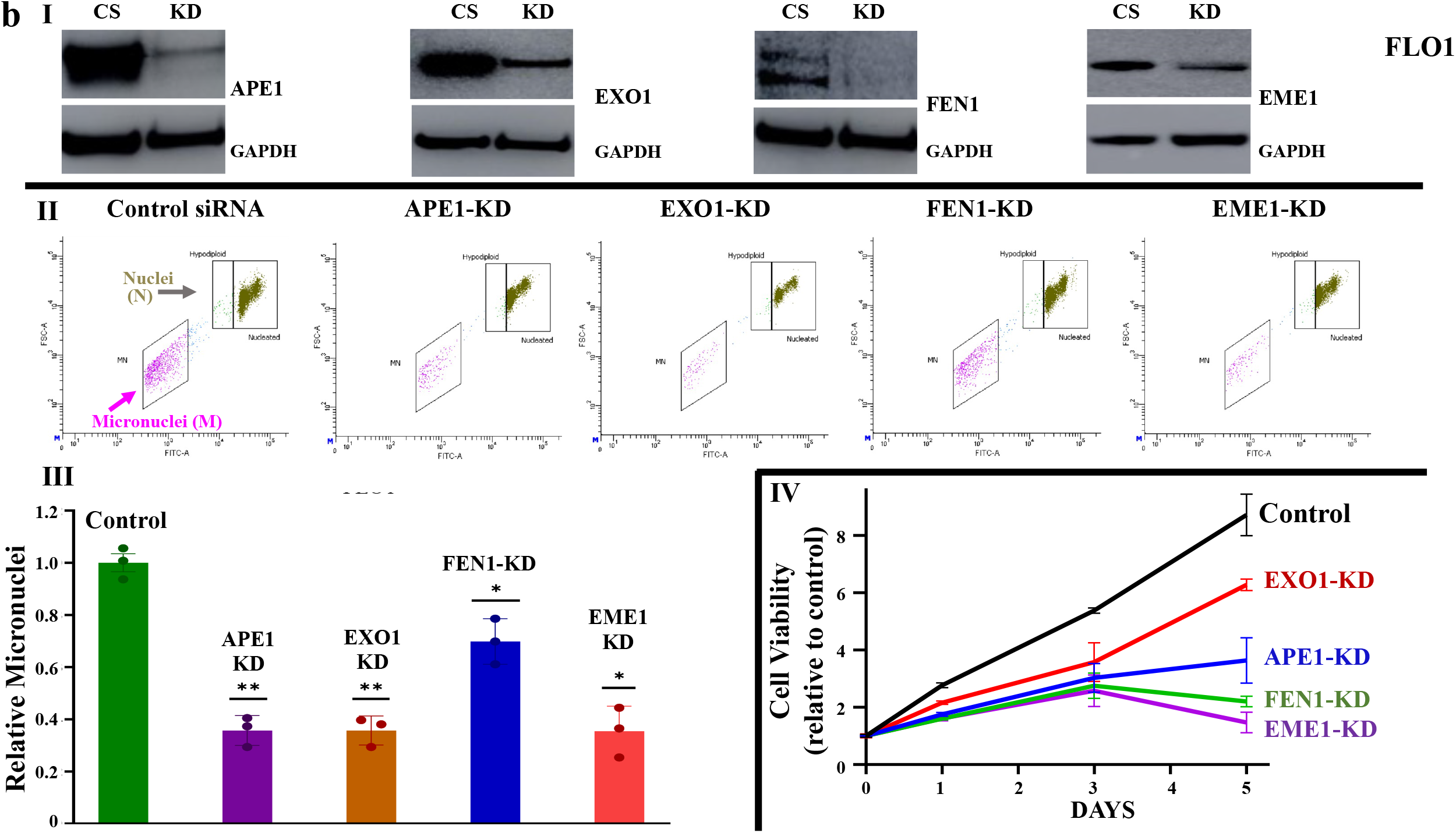
Identification and functional validation of deoxyribonuclease gene signature correlating with genomic instability in solid tumors. **(A) Experimental design, identification of genes and validation plan; (I) Stepwise process leading to identification of genes.** We first assessed gene expression in normal and tumor samples in seven different cancers in TCGA dataset and identified deoxyribonucleases that were overexpressed in these cancers. We then evaluated genomic instability in each patient sample by counting the total number of copy number events; data from triple negative breast cancer (TNBC) patients is shown as an example. Genomic instability data was then integrated with expression data to identify deoxyribonucleases that were overexpressed in cancer relative to corresponding normal samples, and whose expression correlated with genomic instability. We thus identified four gene deoxyribonuclease genomic instability signature (E-GIS) representing six human cancers; **(II) Bar graph showing number of deoxyribonucleases correlating with genomic instability in each cancer.** Four deoxyribonucleases (FEN1, EXO1, EME1 and APE1) correlated with genomic instability in all cancers; **(III) Venn diagram showing genomic instability-associated deoxyribonucleases in six human cancers.** Separate circles for lung adenocarcinoma (LUAD), prostate adenocarcinoma (PRAD), triple negative breast cancer (TNBC) and one common circle for gastrointestinal (GI) cancers i.e., esophageal adenocarcinoma (EAC), stomach adenocarcinoma (STAD) and pancreatic adenocarcinoma (PAAD) are shown. **(IV)** Experimental plan for functional validation of identified genes. **(B) Knockdown screens to validate E-GIS genes for impact on genome stability and growth of cancer cells.** E-GIS genes (APE1, EXO1, FEN1 and EME1) were suppressed in EAC (FLO1) cells using esiRNAs (Millipore/Sigma) and impact on genome stability assessed by evaluating micronuclei (marker of genomic instability) (II) and cell viability (III); CS, control esiRNA; KD, esiRNA-mediated knockdown. **(I)** Western blot showing knockdown of genes; **(II)** Images showing nuclei (N) and micronuclei (MN; a marker of genomic instability). X-axis is FITC signal indicating amount of DNA in nuclei or micronuclei and Y-axis (forward scatter) indicates size which distinguishes nuclei from micronuclei; (**III)** Bar graph showing relative micronuclei; error bars represent SDs of triplicate experiment. Two-tailed p-values derived by Student t-test (*p < 0.05–0.001; **p < 0.001). **(IV)** Cell viability assessed using Cell Titer-Glo; CS, control esiRNA; KD, esiRNA-mediated knockdown; error bars represent SDs of triplicate experiment. Two-tailed p-values derived by Student t-test (p ≤0.03).

Relative to control cells, the knockdown of APE1, EXO1, FEN1 and EME1 in EAC (FLO-1) cells reduced genomic instability by 64%, 64%, 30% and 65% (p ranged from 0.01 to ≤ 0.001), as assessed by micronucleus assay. Impact of knockdown of these genes was also assessed on growth of EAC cells. Relative to control cells, the KD of EXO1, APE1, FEN1 and EME1 was associated with 28%, 58%, 75% and 83% reduction in viable cell number at day 7 after selection (P ≤ 0.03) (Figure 1b, panel IV). Consistently, the overexpression of APEX1, EXO1, FEN1 and EME1 caused 118%, 90%, 45% and 27% increase in genomic instability in FLO-1 cells, respectively (**Supplementary Figure 3, I-II**). Evaluation at day 7 after selection showed that overexpression of APEX1, EXO1, FEN1 and EME1 was associated with 114%, 78%, 42% and 28% increase in viable cell number relative to control cells (Supplementary Figure 3, I and III). Elevated expression of this gene signature also correlated with poor overall survival in pancreatic, lung and one of the esophageal adenocarcinoma datasets (**Supplementary Figure 4**). Overall, these data confirmed the role of N-GIS genes in genomic instability and growth of cancer cells and identified APE1 as the top gene whose suppression or overexpression had the strongest impact on genomic instability in EAC cells. Therefore, APE1 and its inhibitor were investigated in detail for impact on various parameters of genome stability and growth in cancer cell lines representing four different solid cancers.

### APE1 is overexpressed in several solid tumors and contributes to spontaneous and chemotherapy-induced genomic instability

Evaluation in TCGA datasets of six human cancers (EAC, esophageal adenocarcinoma; HPBC, hormone positive breast cancer; TNBC, triple negative breast cancer; LUAD; lung adenocarcinoma; PRAD, prostate adenocarcinoma; COAD, colon adenocarcinoma) demonstrated that relative to corresponding normal samples, APE1 was significantly overexpressed in tumor samples (P < 0.04; Figure 2a). Evaluation of APE1 expression in frozen tissue specimens of normal squamous epithelium (NSE), precancerous Barrett’s esophagus (BE), dysplasia and EAC by immunohistochemistry indicated low expression in normal and precancerous (BE) cells whereas markedly elevated in dysplasia and EAC (Figure 2b, panel I). Significant APE1 expression in EAC vs. normal esophageal tissue specimens was also observed in a tissue array (P < 0.05; Figure 2b, panel II). **Impact on spontaneous genomic instability:** We next evaluated the impact of APE1 on genomic instability in cancer cells. APE1 was suppressed in cancer cell lines (FLO-1 and OE19, esophageal adenocarcinoma; MCF7, breast cancer; A549, epithelial lung carcinoma; PC3, prostate adenocarcinoma) either using lentiviral shRNAs (Figure 2c) or by treatment with APE1 inhibitor (API3) (Figure 2d) and live cell fractions evaluated for micronuclei (a marker of genomic instability) using flow cytometry. Representative images showing nuclei (N) and micronuclei (MN) and bar graphs showing fold change in micronuclei relative to control cells are shown in Figure 2c. Knockdown of APE1 led to a significant reduction (ranging from 45% to 77% reduction; p < 0.05 – < 0.001) in genomic instability in all five cell lines tested (Figure 2c). Consistently, the treatment with APE1 inhibitor in FLO-1, OE19 and MCF7 significantly reduced genomic instability by 68%, 39% and 72%, respectively (p < 0.03; Figure 2d). Moreover, the treatment with APE1 inhibitor in A549 and PC3 reduced genomic instability by ~ 44% in two independent experiments (Figure 2d). **Impact on chemotherapy-induced genomic instability:** Cancer cell lines (OE19, esophageal adenocarcinoma; A549, epithelial lung carcinoma) were treated with APE1 inhibitor (API3), cisplatin (CIS) or combination of both and live cell fractions evaluated for micronuclei. Treatment with cisplatin increased the genomic instability in OE19 and A549 cells by 5.6-fold and 1.6-fold, respectively (Figure 2e). Addition of API3 led to near complete inhibition of cisplatin-induced genomic instability in these cell lines (Figure 2e). These data demonstrate that inhibition of APE1 inhibits spontaneous as well as chemotherapy-induced genomic instability in cancer cells.

**Figure 2.**
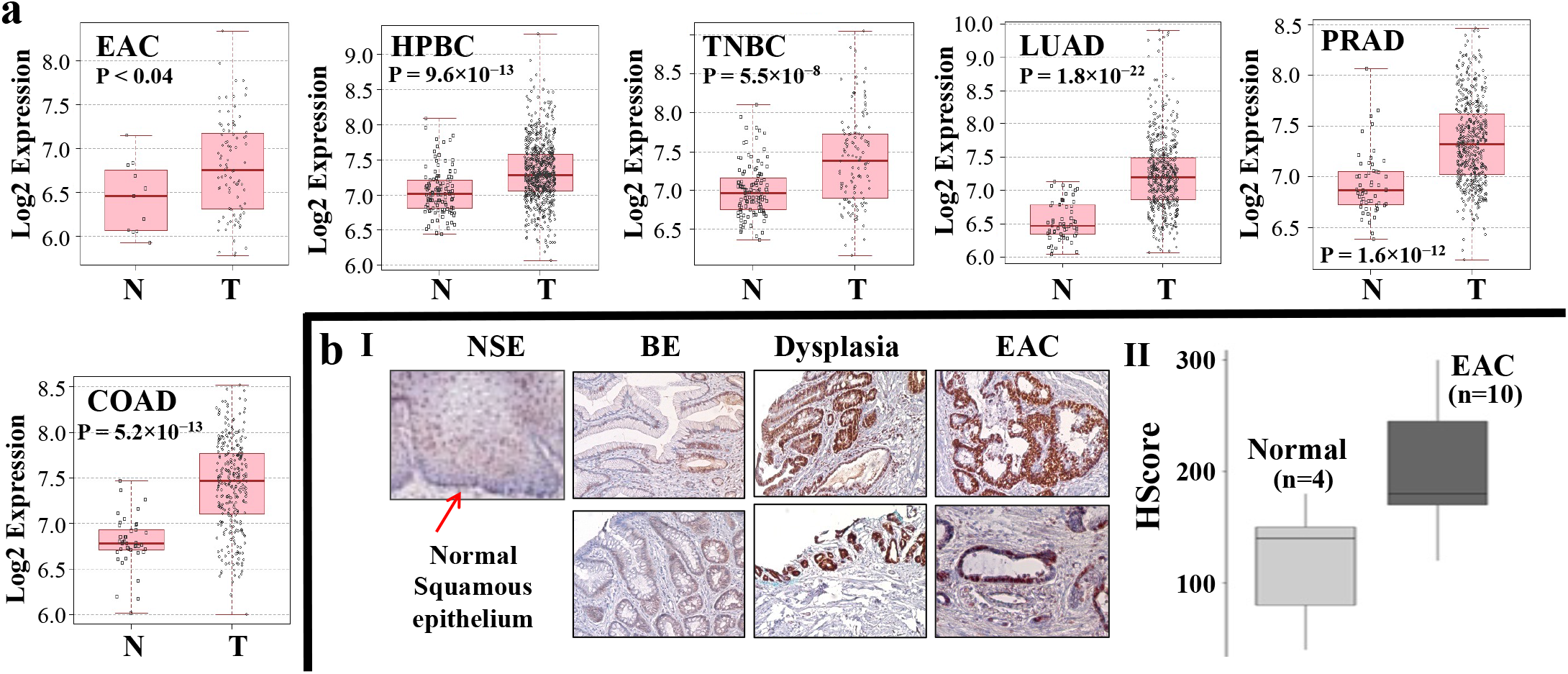
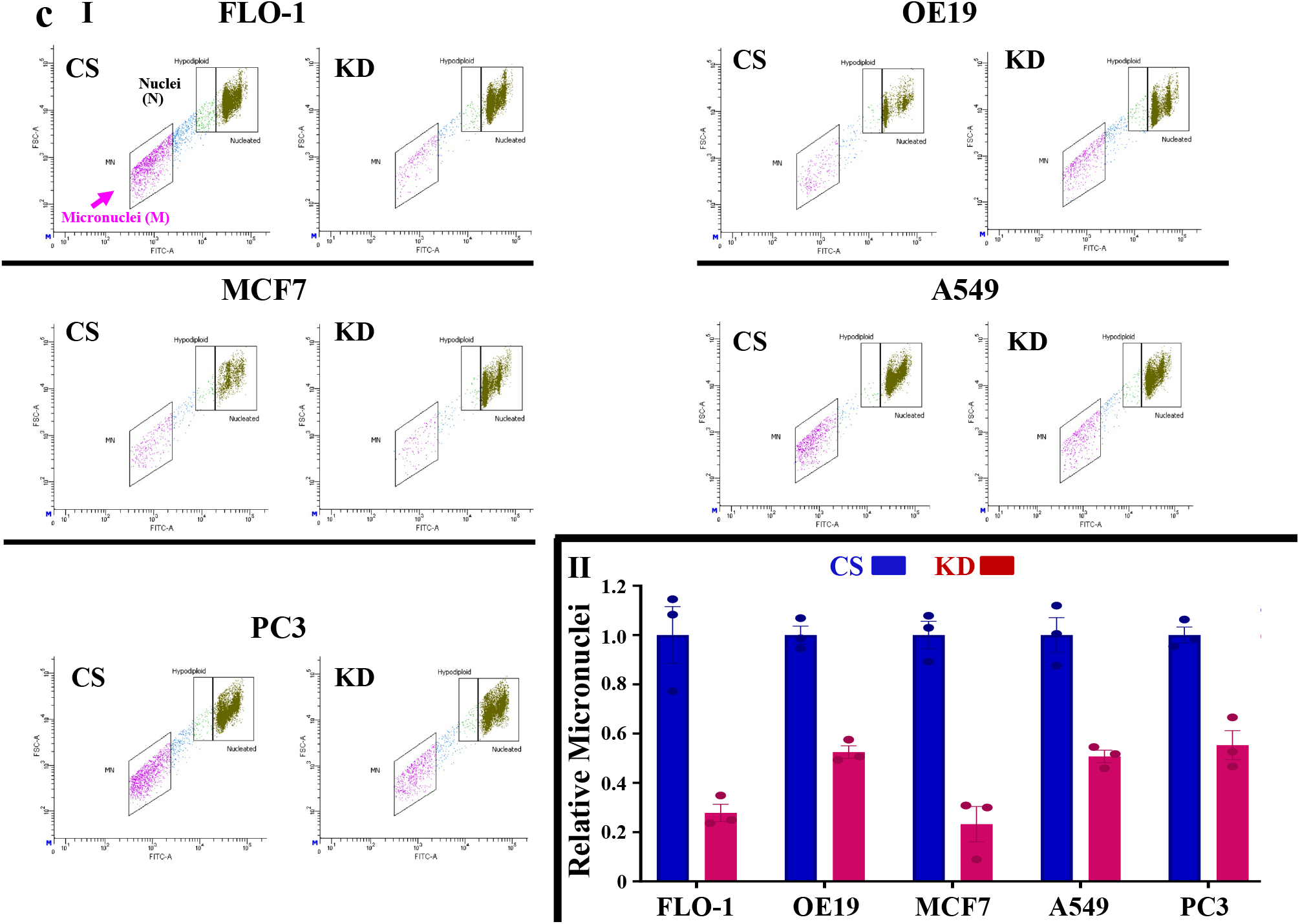
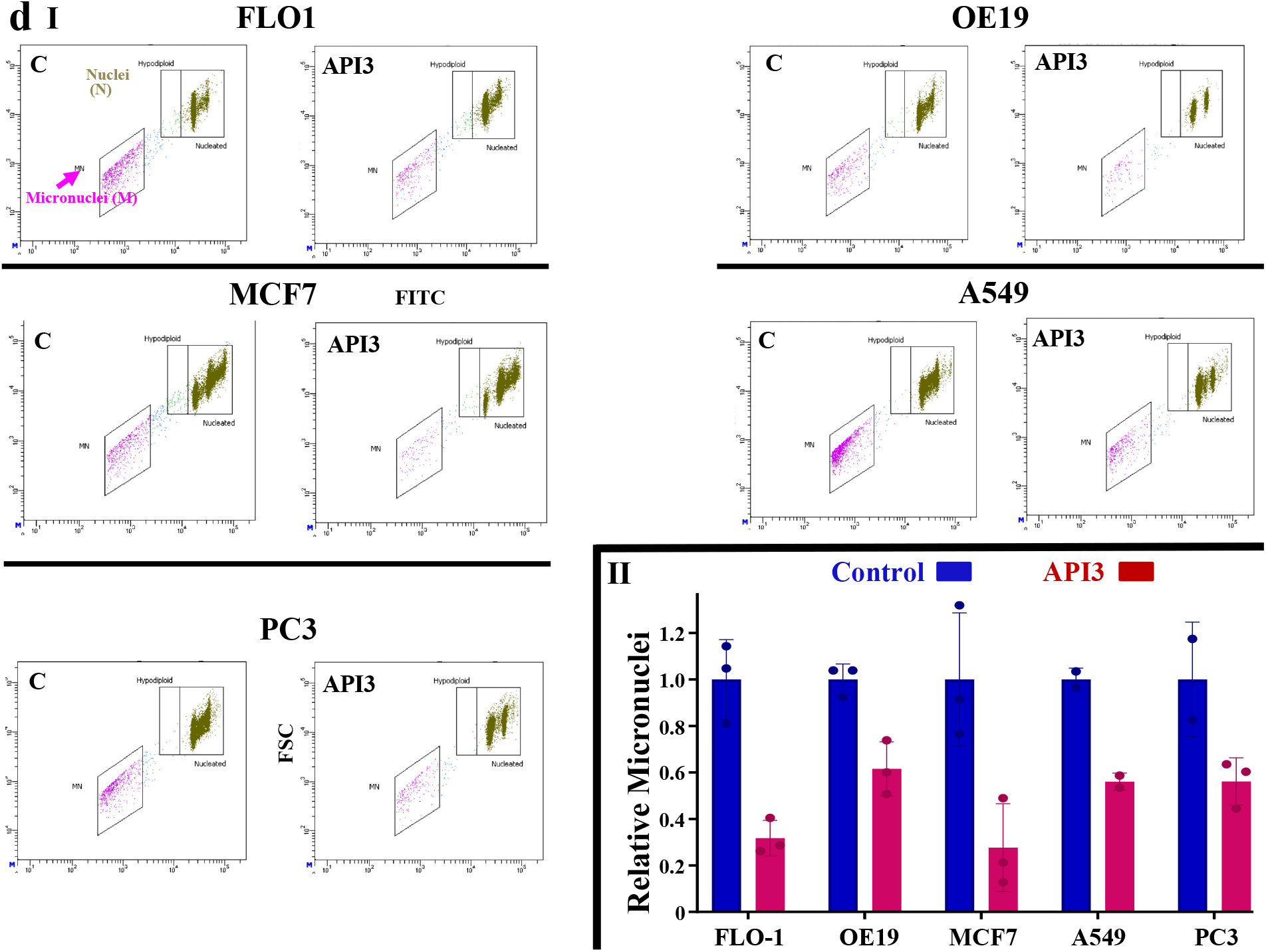
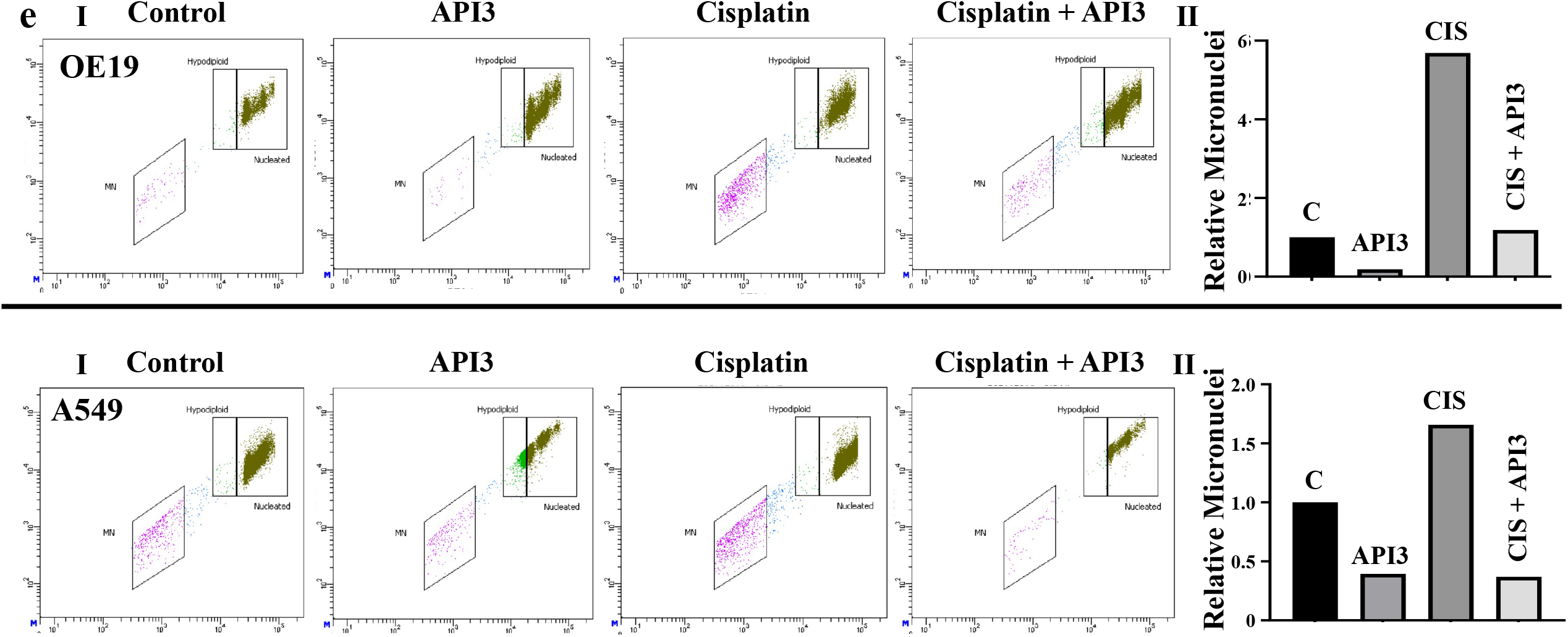
APE1 is overexpressed and contributes to spontaneous as well as chemotherapy-induced genomic instability in EAC and other solid cancer cell lines. **(A-B)** APE1 is overexpressed in solid tumors. Relative expression (Log2) of APE1 in TCGA datasets was plotted using R and normal and tumor samples compared with limma software. EAC, esophageal adenocarcinoma; HPBC, hormone positive breast cancer; TNBC, triple negative breast cancer; LUAD; lung adenocarcinoma; PRAD, prostate adenocarcinoma; COAD, colon adenocarcinoma. **(B)** APE1 is overexpressed in EAC patient samples; Panels: (I) APEX1 expression in frozen tissue specimens of normal squamous epithelium (NSE), Barrett’s esophagus (BE), dysplasia and EAC detected by immunohistochemistry; (II) APEX1 expression in EAC and normal esophageal tissue specimens on a tissue array (Biomax); **(C-E)** Impact on genomic instability: **(C)** APE1 was suppressed in cancer cell lines (FLO-1 and OE19, esophageal adenocarcinoma; MCF7, breast cancer; A549, epithelial lung carcinoma; PC3, prostate adenocarcinoma) using lentiviral shRNAs and right after selection, live cell fractions evaluated for micronuclei (marker of genomic instability) using flow cytometry. Panels: (I) Representative images showing nuclei (N) and micronuclei (MN). X-axis is FITC signal indicating amount of DNA in nuclei or micronuclei and Y-axis (FSC or forward scatter) indicates size which distinguishes nuclei from micronuclei; (II) Bar graph showing relative micronuclei in control and knockdown cells; Error bars represent SDs of four independent experiments. Two-tailed p-values for significance of difference between control and each treated sample was derived by Welch’s t-test (*p < 0.05–0.001; **p < 0.001). CS, control shRNA; KD, cells transduced with APE1 shRNA; **(D)** All five cell lines (described above) were treated with API3 (1.5 ⍰M) for 48 h and live cell fractions evaluated for micronuclei (marker of genomic instability) using flow cytometry. Panels: (I) Representative images showing nuclei (N) and micronuclei (MN); (II) Bar graph showing relative micronuclei in control and treated cells; C, control (DMSO) treated cells; API3, API3-treated cells; For FLO-1, OE19 and MCF7 error bars represent SDs of four independent experiments and two-tailed p-values for significance of difference between control and each treated sample derived by Welch’s t-test (p < 0.03). Bar graphs for A549 and PC3 cell lines represent two independent experiments; **(E)** Cancer cell lines, esophageal adenocarcinoma (OE19. top panel) and epithelial lung carcinoma (A549; bottom panel) were treated with APE1 inhibitor (API3; 1.5 ⍰M), cisplatin (CIS; 5 ⍰M) or combination of both and live cell fractions evaluated for micronuclei (marker of genomic instability) using flow cytometry. Panels: (I) Representative images showing nuclei (N) and micronuclei (MN); (II) Bar graph showing fold change in micronuclei, relative to control cells.

### APE1 contributes to increased DNA breaks, RAD51 expression and homologous recombination activity. Impact on cisplatin-induced DNA breaks

APE1 was inhibited in cancer cell lines (FLO-1, esophageal adenocarcinoma; A549, epithelial lung carcinoma; MCF7, breast cancer) using an inhibitor (API3) or gene knockdown. Control and APE1-inhibited cells were treated with cisplatin and live cell fractions evaluated for expression of APE1 and **γ**-H2AX (marker of DNA breaks) by Western blotting. Treatment with cisplatin led to a substantial increase in DNA breaks whereas chemical as well as transgenic suppression of APE1 inhibited cisplatin-induced DNA breaks in all five cancer cell lines (Figure 3a, I-II). **Impact on homologous recombination (HR) activity:** Our previous data in multiple myeloma demonstrated that APE1 regulates HR activity^51^, an important mechanism of genomic instability in cancer^11,12^. We, therefore, investigated the impact of APE1 inhibition on HR in solid tumor cells. Treatment with API3 inhibited RAD51 promoter activity (Figure 3b - I), RAD51 expression (Figure 3b - II) and HR activity (Figure 3b - III) in EAC ( FLO-1) cells. Consistent with chemical inhibition, shRNA-mediated knockdown of APE1 (shown Figure 3c - I) also inhibited RAD51 expression (Figure 3c - II), its phosphorylation (Figure 3c - III) and HR activity (Figure 3c - IV) in EAC cells. Treatment of other cancer cell lines (OE19, esophageal adenocarcinoma; MCF7, breast cancer; A549, epithelial lung carcinoma; PC3, prostate adenocarcinoma) representing four solid cancer types with APE1 inhibitor (API3) also resulted in a dose-dependent inhibition of HR activity (Figure 3d; p < 0.05). Consistently, the suppression of APE1 in these cell lines using shRNA also led to a significant inhibition of HR activity (p values ranged from < 0.05 to < 0.0001; **Supplementary Figure 5**). To further understand the functional link between APE1 and HR, we identified APE1-interacting proteins in EAC (FLO-1) cells by mass spectrometry. Top APE1 interactors were selected by protein-protein network analysis (using String) and their pathways monitored. The proteins interacting with APE1 belonged to several DNA repair pathways including those related to HR. These include double-strand break repair, DNA recombination, DNA duplex unwinding, response to gamma radiation, telomere maintenance, base excision repair and non-homologous end joining. Several proteins with known functional role in HR including RPA1^59^, WRN^60^, DHX9^61^, ILF2^62^ and YBX1^62^ were among top interactors (**Supplementary Figure 6)**. These data also support the functional link between APE1 and HR pathway. Overall, these data demonstrate that elevated APE1 contributes to increased DNA breaks and HR activity in EAC as well as other solid tumors.

**Figure 3.**
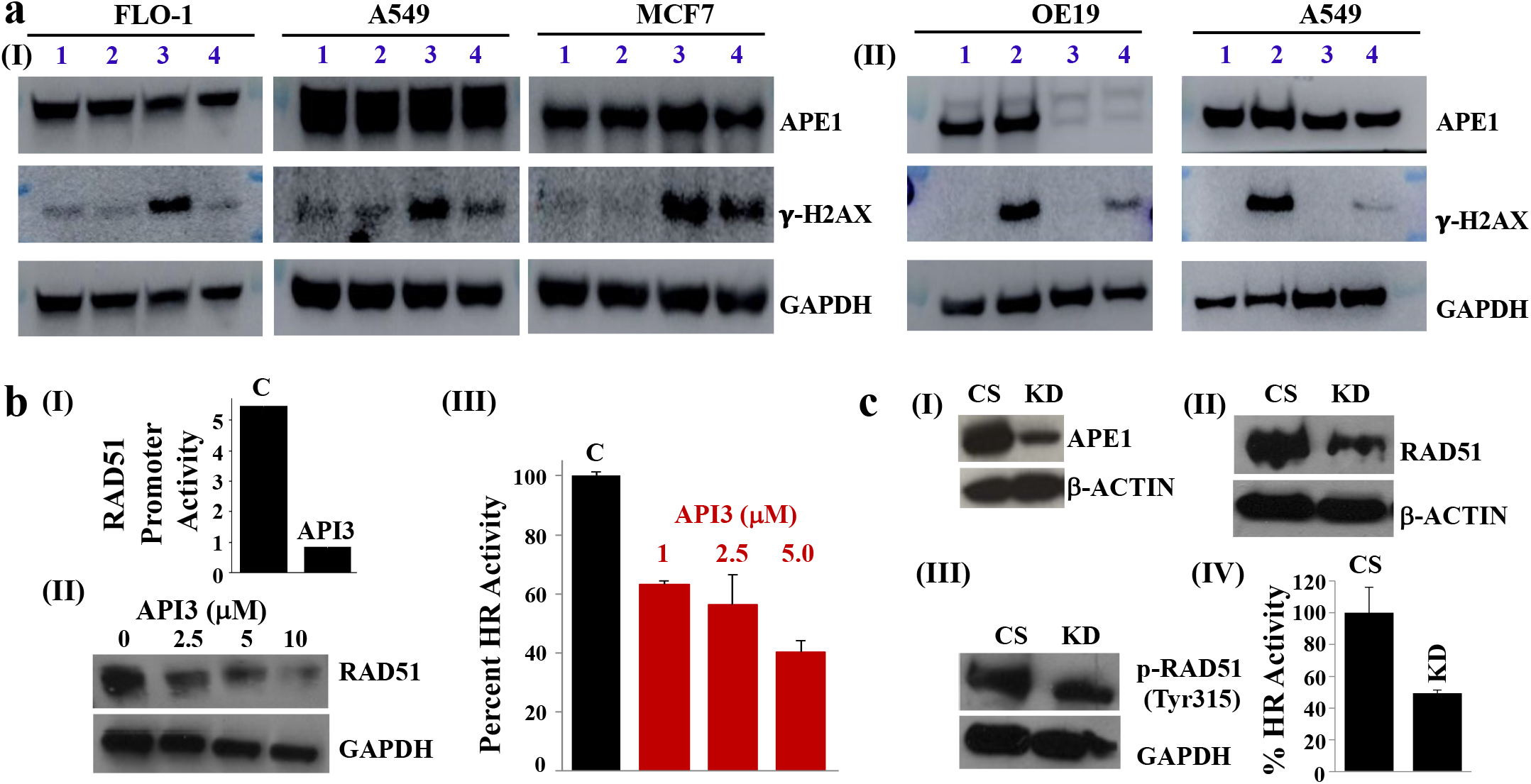
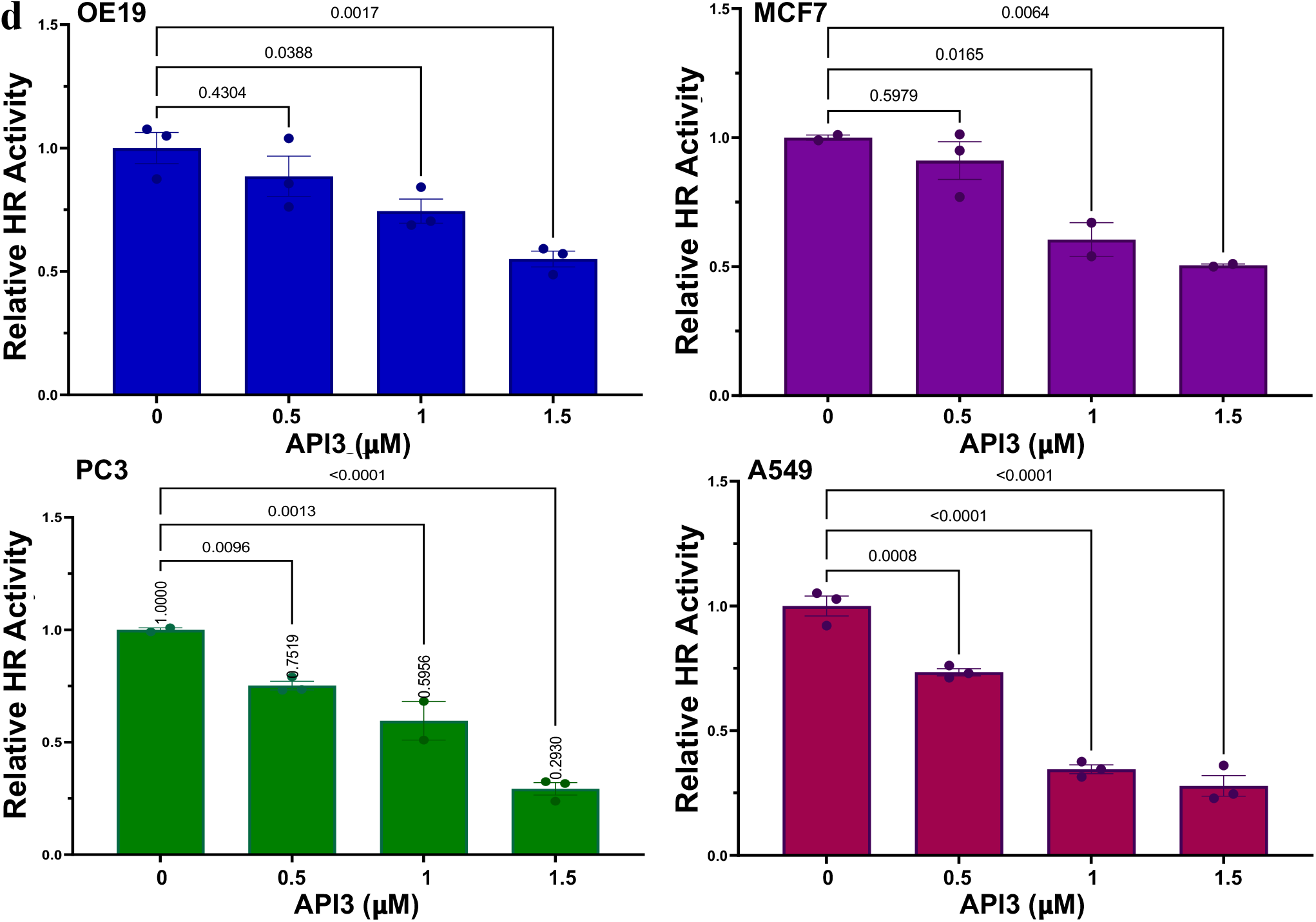
APE1 contributes to increased DNA breaks, RAD51 expression and homologous recombination (HR) activity in cancer cells. **(A) Chemical or transgenic suppression of APE1 reduces cisplatin-induced DNA breaks in solid cancer cells. (I)** Cancer cell lines (FLO-1, esophageal adenocarcinoma; A549, epithelial lung carcinoma; MCF7, breast cancer) were treated with APE1 inhibitor (API3; 1.5 ⍰M), cisplatin (5 ⍰M) or combination of both and live cell fractions evaluated for expression of APE1 and **γ**-H2AX (marker of DNA breaks) by Western blotting. Lanes: 1, control; 2, API3; 3, cisplatin; 4, API3+cisplatin. **(II)** Control and APE1-knockdown cancer cells (OE19, esophageal adenocarcinoma; A549, epithelial lung carcinoma) were treated with cisplatin (15 ⍰M) and expression of APE1 and **γ**-H2AX assessed by Western blotting. Lanes: 1, control shRNA; 2, control shRNA treated with cisplatin; 3, APE1-shRNA; 4, APE1-shRNA treated with cisplatin; **(B-C) Chemical or transgenic inhibition of APE1 inhibits RAD51 expression and HR activity. (B)** EAC ( FLO-1) cells treated with API3 were evaluated for RAD51 expression by Western blotting (I), RAD51 promoter activity using a plasmid in which RAD51 drives luciferase expression (II), and HR activity using the plasmid-based assay (III). **(C)** EAC (FLO-1) cells were treated with shRNAs (CS, control shRNAs; KD, cells treated with APE1-targeting shRNAs). Panels: (I) APE1 expression evaluated by Western blotting; (II) HR activity assessed by plasmid-based assay; (III) Expression of RAD51 (top panel) and its phosphorylated form (bottom panel) evaluated by Western blotting; **(D) Dose-dependent inhibition of HR activity by APE1 inhibitor in solid tumor cell lines.** Cancer cell lines (OE19, esophageal adenocarcinoma; MCF7, breast cancer; A549, epithelial lung carcinoma; PC3, prostate adenocarcinoma) were treated with APE1 inhibitor (API3) at different concentrations for 48 HR and impact of HR activity determined using plasmid-based functional assay described in Methods.

### APE1 inhibition impairs growth and increases cytotoxicity of chemotherapeutic agent *in vitro* and *in vivo*. Impact on growth and colony formation

APE1 was suppressed in human cancer cell lines-esophageal adenocarcinoma (FLO-1, OE19), breast cancer (MCF7), epithelial lung carcinoma (A549) and prostate adenocarcinoma (PC3) using shRNAs and impact on cell viability assessed using Cell Titer-Glo. Relative to control shRNA, APE1-knockdown (KD) reduced the growth rate of all five cancer cell lines. Evaluation at day 7 after selection revealed that relative to control cells, the knockdown of APE1 in FLO1, OE19, MCF7, A549 and PC3 cells was associated with reduction in cell viability by 47%, 70%, 56%, 43% and 76%, respectively (P < 0.02) (**Figure 4a**). APE1 inhibition, by inhibitor or gene knockdown, strongly impaired colony formation in all five cell lines (**Supplementary Figure 7**).

**Figure 4.**
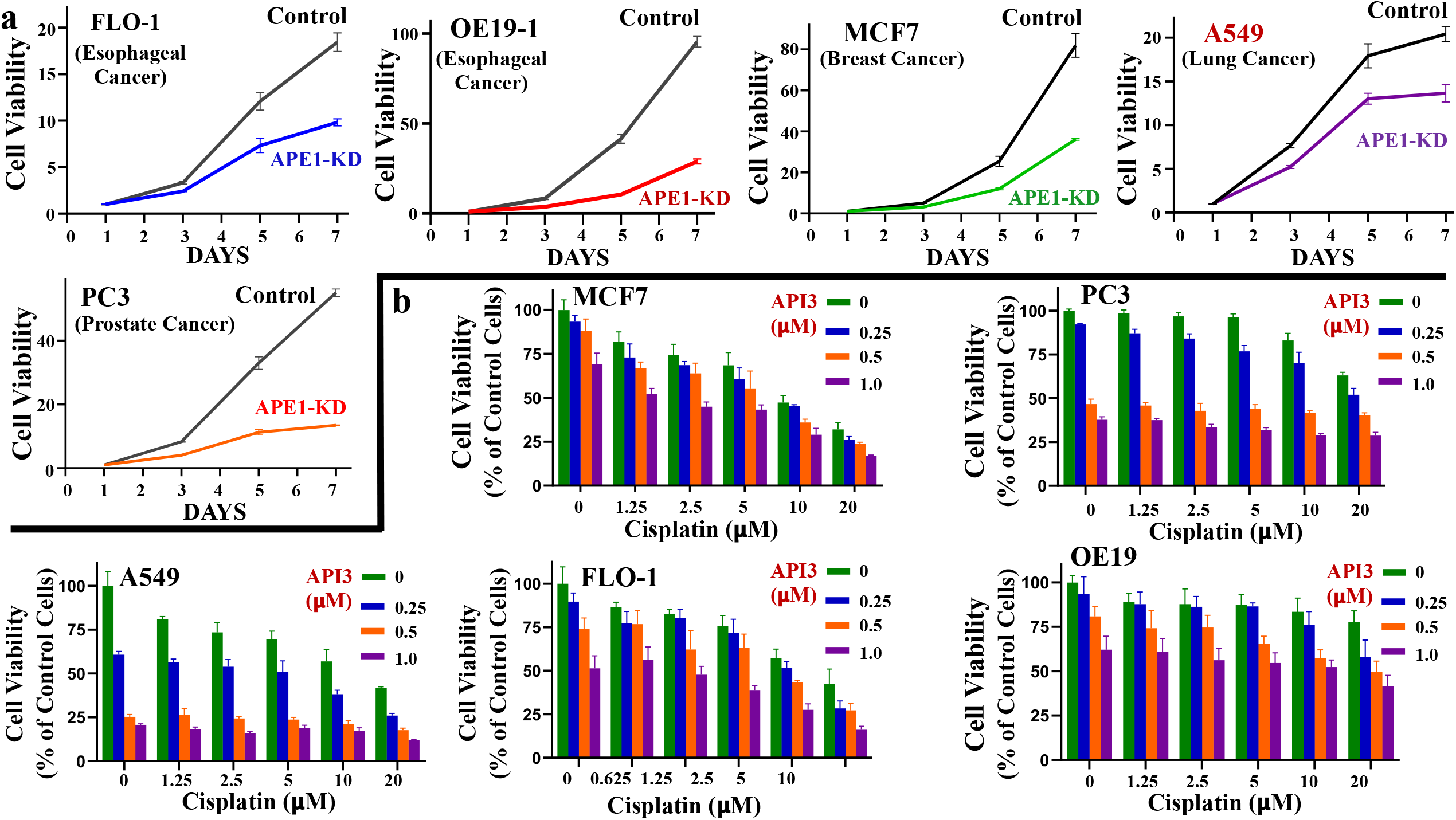
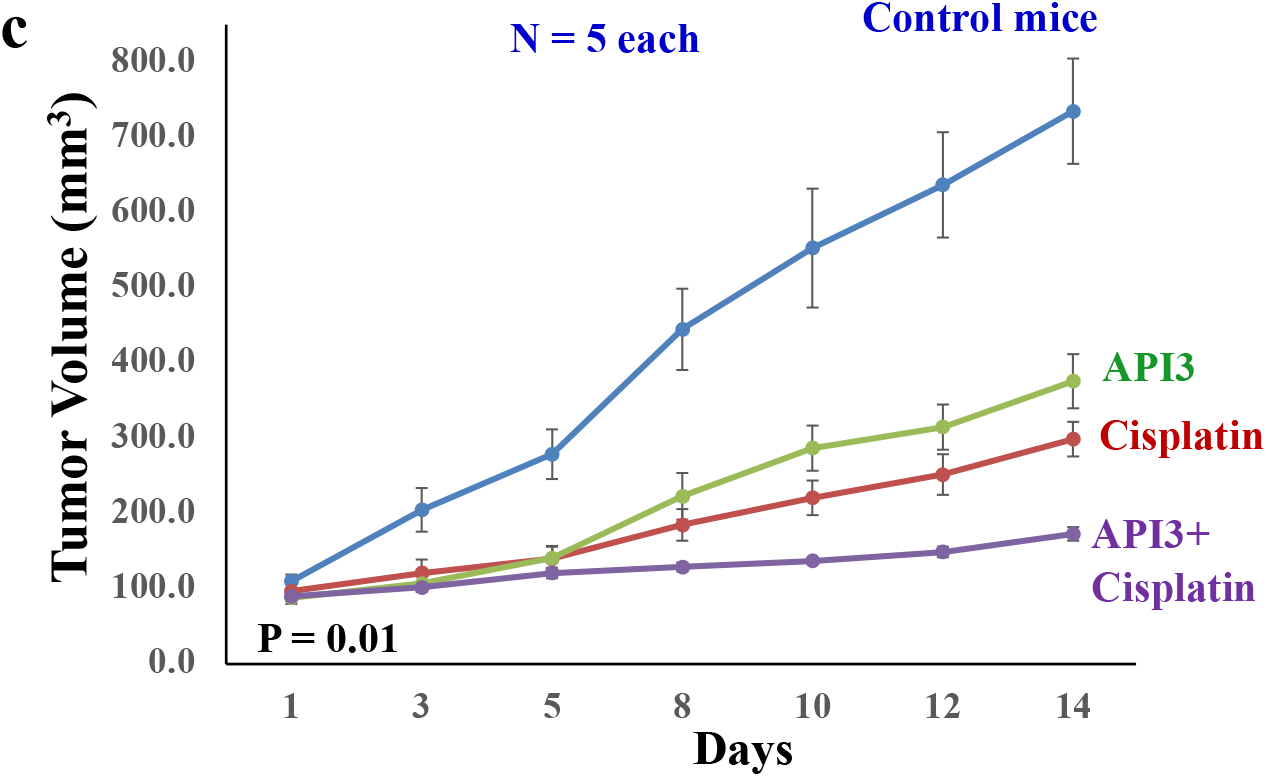
Suppression of APE1 inhibits growth and increases cytotoxicity of chemotherapeutic agent in cancer cells *in vitro* and *in vivo*. **(A) APE1-knockdown inhibits growth of solid cancer cell lines.** APE1 was suppressed in human cancer cell lines - esophageal adenocarcinoma (FLO-1, OE19), breast cancer (MCF7), epithelial lung carcinoma (A549) and prostate adenocarcinoma (PC3) using shRNAs and impact on cell viability assessed using Cell Titer-Glo; Control, control shRNA; APE1-KD (knockdown), APE1 shRNA; error bars indicate SDs of triplicate experiment. Two-tailed p values for significance of difference between control and knockdown cells at day 7 ranged from 0.015 to 0.0007. **(B) APE1 inhibitor increases cytotoxicity of cisplatin *in vitro*.** Cancer cell lines (MCF7, A549 and FLO-1) were treated with APE1 inhibitor (API3), alone as well as in the presence of chemotherapeutic agent cisplatin, and cell viability measured after 48 hr. Bar graphs of cells treated with API3 with cisplatin are shown; error bars represent SDs of three experiments. Combination index plots (calculated using CalcuSyn software) are shown in Supplementary Figure 8. **(C) APE1 inhibitor inhibits cancer cell growth and increases cytotoxicity of cisplatin *in vivo*.** EAC (OE19) cells were injected subcutaneously in SCID mice and following appearance of tumors, mice treated with either DMSO, API3 (12 mg/kg, daily for 2 weeks), cisplatin (3 mg/kg, once a week for 2 weeks) or combination of both drugs. Line plots showing tumor growth in control and treated mice (n = 5 each); error bars represent SDs.

### Impact on cytotoxicity of chemotherapeutic agent

Cancer cell lines (MCF7, A549, FLO-1, OE19 and PC3) were cultured in the presence of APE1 inhibitor (API3), alone or in combination with cisplatin for 48 h, and cell viability assessed. API3 treatment led to increased cytotoxicity of cisplatin in cancer cells (Figure 4b); combination index plots show that increase in cytotoxicity by combination treatment was mostly synergistic or additive in all cancer cell lines tested **(Supplementary Figure 8)**. **Impact on tumor growth and cytotoxicity of cisplatin in vivo.** To evaluate the efficacy of APE1 inhibitor (API3), alone and in combination with a chemotherapeutic agent, EAC (OE19) cells were injected subcutaneously in SCID mice and following the appearance of tumors, mice treated with vehicle control, API3, cisplatin or combination of both. Relative to average tumor size in control mice, the tumor size in mice treated with API3, cisplatin and combination reduced by 48.8% (p = 0.0018), 59.3% (p =0.00035) and 76.5% (p = 4.49E-05), respectively (Figure 4c). Average tumor volume in mice treated with combination of both drugs was significantly smaller than treatment with either drug alone (p < 0.001) (Figure 4c). Overall, these data show that APE1 inhibitor impairs cancer cell growth both in vitro and in vivo and synergistically increases the efficacy of chemotherapeutic agents.

### APE1 overexpression in normal cells dysregulates DNA repair and genome stability

Normal primary human esophageal epithelial cells (HEsEpiC; ScienCell) were transfected with control plasmid (C) or the plasmid carrying APE1 gene under a strong promoter to overexpress APE1 (APE1O). **DNA breaks and homologous recombination (HR):** Upregulation of APE1 (shown in Figure 5a I) was associated with increased DNA breaks, as assessed from **γ**-H2AX expression (Figure 5A I) as well as by Comet assay (Figure 5A II). APE1 overexpression also led to increased RAD51 expression (Figure 5A I), HR activity (Figure 5A III) and chromosomal instability as evident from centrosome amplification (Figure 5B). **Karyotypic instability:** To further investigate the impact of APE1 upregulation on chromosomal instability, the cells were cultured for 60 days, and their chromosomes visualized by spectral karyotyping. The control cells had near diploid karyotype with mitotic index (MI) of 0.16% and had 0.2 aberrations/cell (Figure 5C I), whereas APE1-overexpressing (APE1O) cells were near tetraploid with MI of 3.9%, 16 aberrations/cell, 4 copies of C-myc/cell and several chromosomal abnormalities, indicating a striking karyotypic instability (Figure 5C II-III). **Whole genome sequencing demonstrates mutational instability:** Genomic impact of APE1-overexpression in normal cells was evaluated by whole genome sequencing. Genome of day 0 cells was used as baseline to identify the mutations acquired by control and APE1-overexpressing cells over a period of 60 days in culture. Although, the cells carrying control plasmid had only 83 new mutations, those acquired by APE1-overexpressing cells were > 25,000 (Figure 5D I). Removal of known SNPs from these data identified 3500 mutations unique to APE1-overexpression. These mutations were further investigated for subtypes and signatures of underlying mutational processes as described by us previously^10,54^. Most of the mutations caused by APEI-overexpression were C>T substitutions which were followed by T>C and then C>A (Figure 5D II). Investigation of mutational signatures identified Signature 3, indicative of HR dysfunction, as the top mutational process activated by APE1 (Figure 5D III). Second most frequent signature was Signature 1. This signature reflects a mutational process started by spontaneous deamination of 5-methylcytosine which could be the basis for C > T mutations.

**Figure 5.**
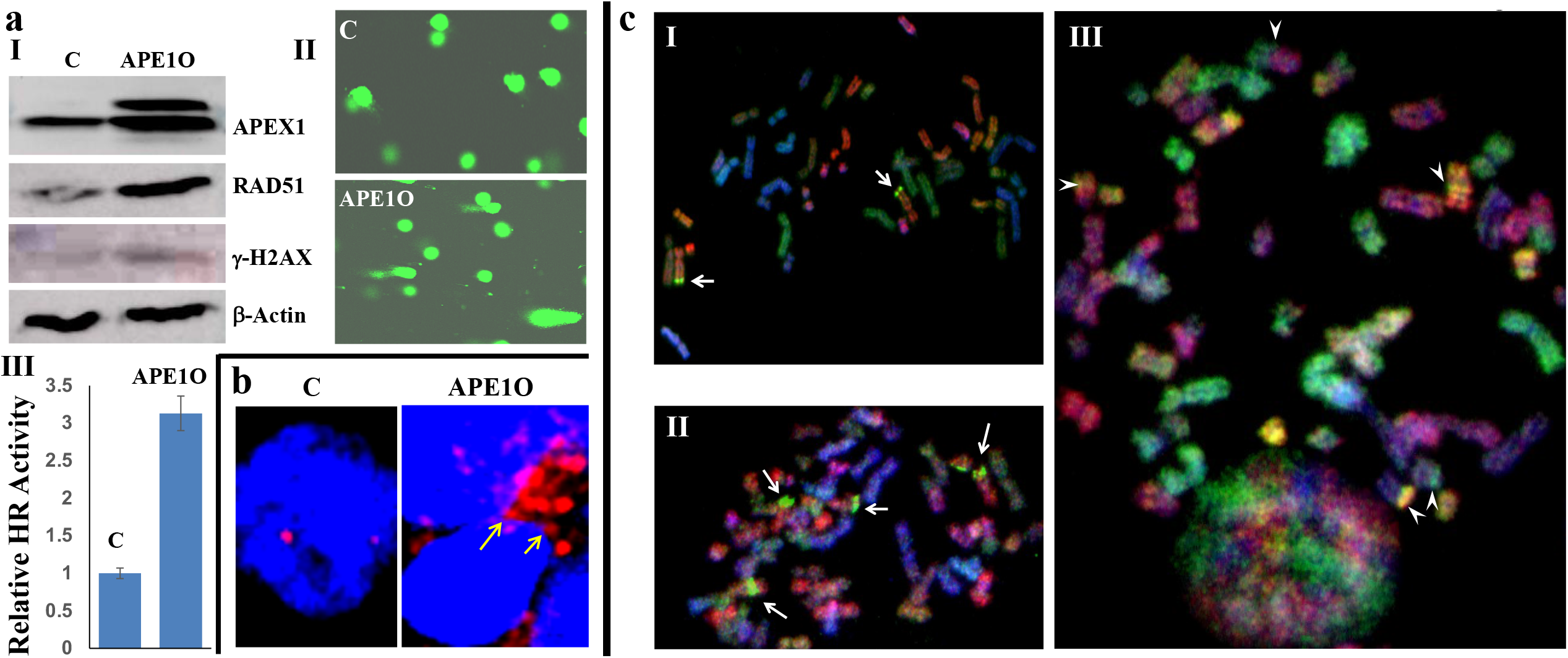
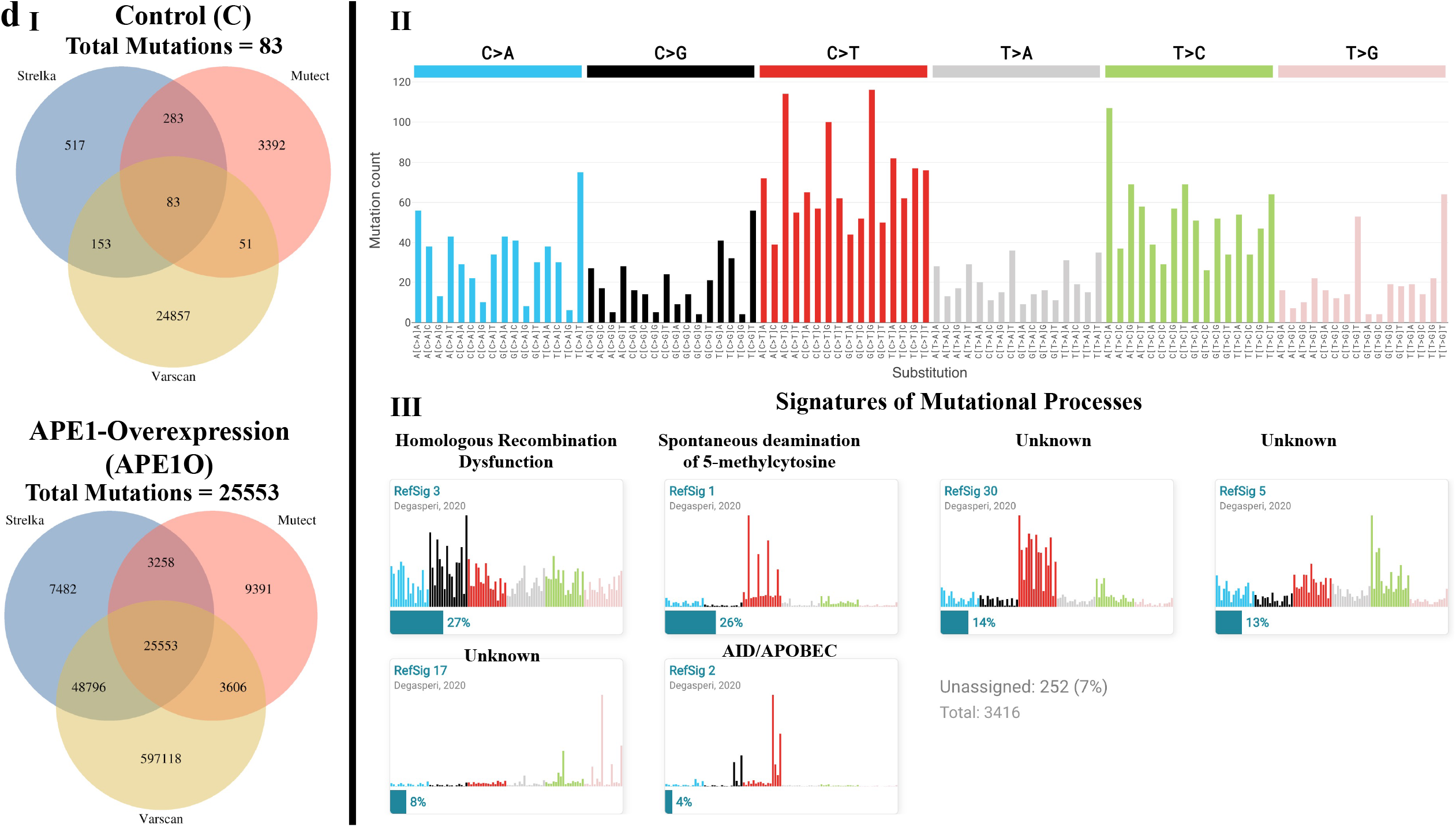
Overexpression of APE1 in normal esophageal cells induces DNA breaks, homologous recombination and chromosomal and mutational instability. Normal primary human esophageal epithelial cells (HEsEpi; ScienCell) were transfected with plasmids carrying empty control vector (C) or APE1 under a strong promoter (APE1O) and evaluated for various parameters of genome stability. **(A) DNA breaks and homologous recombination (HR). (I)** Western blot showing expression of APE1, RAD51 and γ-H2AX (marker of DNA breaks) in control and APE1-overexpressing cells, evaluated right after selection; **(II)** DNA breaks assessed by Comet assay right after selection; **(III)** HR activity evaluated by plasmid-based assay; error bars represent SD of three experiments and two tailed p value is < 0.0002; **(B) Impact on centrosome:** Centrosome investigated three weeks after selection. Arrows show multiple centrosomes clustered together in APE1O cells; **(C) Impact on karyotype:** Karyotypes of control (**I**) and APE1O (**II-III**) cells examined after 60 days in culture. Panels II and III are two different examples of karyotypes. Mitotic index (MI) of APE1O = 3.9%; 9-12 chromosome aberrations (arrowheads on representative translocations); **(D) Impact on mutational frequency evaluated by WGS:** Normal primary esophageal epithelial cells carrying control plasmid (C) and that for APE1-overexpression (APE1O) were cultured for sixty days and analyzed for new mutations, relative to baseline (day 0) cells, using WGS. **(I)** New mutations in control and APE1O cells identified by 3 different software tools; **(II)** Substitution types in APE1-overexpressing cells throughout genome are presented in context of the sequence immediately 5’ and 3’ to the mutated base; **(III)** Fraction of contribution of each mutation type at each sequence context for the mutational signatures. For investigation of mutational signatures or processes, SNPs were removed to ensure that somatic mutation callers do not confuse about the SNPs and SNVs. After removing SNPs, there were 44 mutations in control vs. 3500 mutations in APE1-overexpression cells.

DNA samples from control and APE1-overexpressing cells cultured for a period of 60 days were also investigated for copy number alterations using single nucleotide polymorphism arrays. Relative to day 0 cells, the copy number events acquired in control and APE1-overexpressing cells were 66 and 5734, respectively (**Supplementary Figure 9**). These data are commensurate with that observed by whole genome sequencing.

### APE1-induced genomic instability can predispose normal cells to oncogenic transformation

Consistent with increased DNA and karyotypic instability, the APE1-overexpressing cells changed in morphology from elongated to somewhat rounded in shape (Figure 6A) and had significantly increased growth rate relative to control plasmid-transfected cells (Figure 6B). **Tumors in xenograft model:** When injected subcutaneously in SCID mice, control plasmid-transfected cells (injected on left side of each mouse) did not form tumors, but APE1-overexpressing cells (injected right side of each mouse) formed tumors (Figure 6C). These data demonstrate that elevated APE1 activity drives genomic instability, predisposing normal cells to tumorigenesis. **Karyotypes of mice tumors:** Tumors from euthanized mice were removed and investigated for chromosomal aberrations as described in methods. The karyotypes were described relative to a diploid status indicated in the karyotype by < 2n > since the parent line was pseudodiploid. This is to be noted that the distinctive aberrations seen in the parent line were not identifiable in this xenograft, presumably due to extensive further rearrangement. All metaphases observed were aneuploid and highly abnormal. Chromosome count ranged from 54-68 (hyperdiploid to hypertriploid) in the 20 cells examined, with a modal number of 58 (5/20 cells). Chromosome abnormalities that were clonal (seen in at least two karyotypes for chromosome gains and in at least three karyotypes for loss) included gains as well as loss of entire chromosomes (Figure 6D; **Supplementary Figure 10**). In some cases, gains of partial regions of chromosomes were evident, due to additional copies of structurally abnormal chromosomes. Structural chromosome abnormalities, many of them only partially identifiable with regard to chromosomal origin, were seen in every karyotype. Clonal abnormalities, occurring in at least some karyotypes, are indicated by solid arrows (Figure 6D; Supplementary Figure 10). Six clonal markers, with distinctive banding patterns but whose chromosomal origin is unknown were identified, mar1-mar6. In addition, all karyotypes had apparently non-clonal structural abnormalities, placed in the karyotype if partially identifiable, and 7-11 unidentifiable marker chromosomes (placed below the numbered chromosomes), whose clonality could not be established by banding. Overall, this complex and highly aneuploid karyotype suggests ongoing genetic instability.

**Figure 6.**
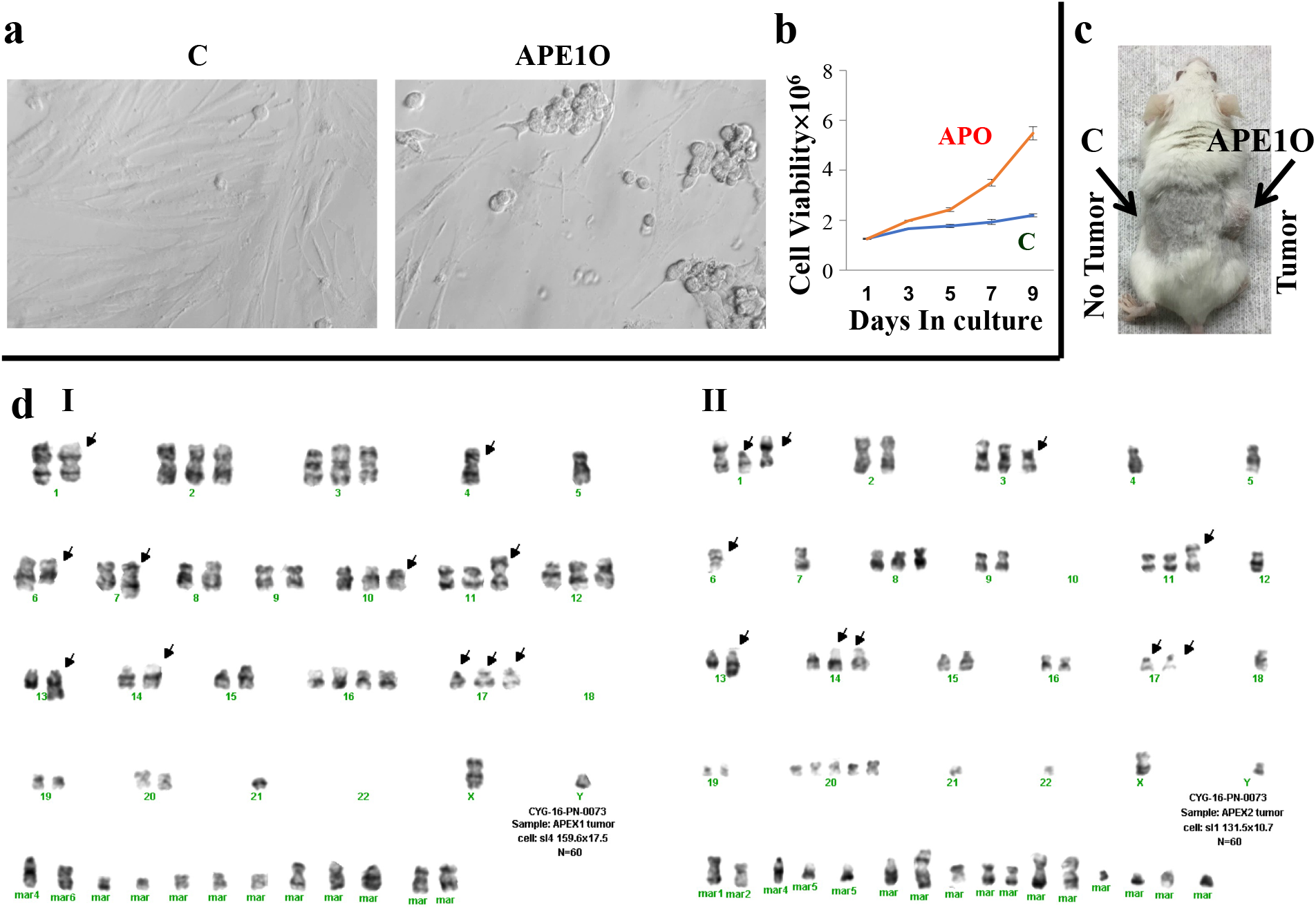
APE1 overexpression leads to oncogenic transformation of normal human esophageal epithelial cells. **(A)** Control and APE1O (APEX1-overexpressing) human esophageal epithelial cells photographed under a phase contrast microscope; **(B)** Growth rate of control and APE1O cells. Cells were cultured and viable cell number assessed at indicated time points using Cell Titer-Glo; error bars represent SDs of triplicate experiment; **(C)** Control and APE1O human esophageal epithelial cells were subcutaneously injected in SCID mice and evaluated for tumor formation. Control cells injected on left side of mice did not grow as tumor whereas APE1O cells injected on right side grew as tumor; one mouse is shown as an example; **(D)** Tumors from control and APE1O mice were evaluated for karyotype. Tissue was minced, dissociated and cells cultured. Metaphases obtained at day 4 were used for analysis; Panels I and II are two representative karyotypes.

### APE1 contributes to the regulation of G2/M checkpoint. Impact on G2/M checkpoint

To investigate the role of APE1 in solid tumors, we either suppressed it in EAC (FLO-1) cells or overexpressed it in normal esophageal epithelial cells and evaluated the impact on expression profile in these cells. Investigation of the pathways that were downregulated in EAC cells treated with APE1 inhibitor whereas upregulated following APE1-overexpression in normal cells, identified cell cycle G2/M checkpoint as the most significant common pathway impacted (**Figure 7a I**). **Impact on G2/M checkpoint in EAC cells:** To further evaluate the role of APE1 in the regulation of G2/M checkpoint in EAC cells, FLO-1 cells were treated with API3 (2 ⍰M) for 48 h and impact on cell cycle evaluated by flow cytometry (Figure 7a, II-IV). Relative to control cells, the treatment with APE1 inhibitor led to 2.4-fold increase in G2 phase of cell cycle (p = 1.5×10^-5^) (Figure 7a, IV). **Impact on G2/M checkpoint in different cancers.** Human cancer cell lines-esophageal adenocarcinoma (OE19), breast cancer (MCF7), epithelial lung carcinoma (A549) and prostate adenocarcinoma (PC3) were treated with API3 for 48 h and evaluated for impact on cell cycle. APE1 inhibition increased the percentage of cells in G2 phase of cell cycle (ranging from ~ 20% to 80% increase) in all these cell lines (Figure 7b). In summary, we show that APE1 is involved in the regulation of cell cycle in cancer cells and its inhibition causes G2/M arrest.

**Figure 7.**
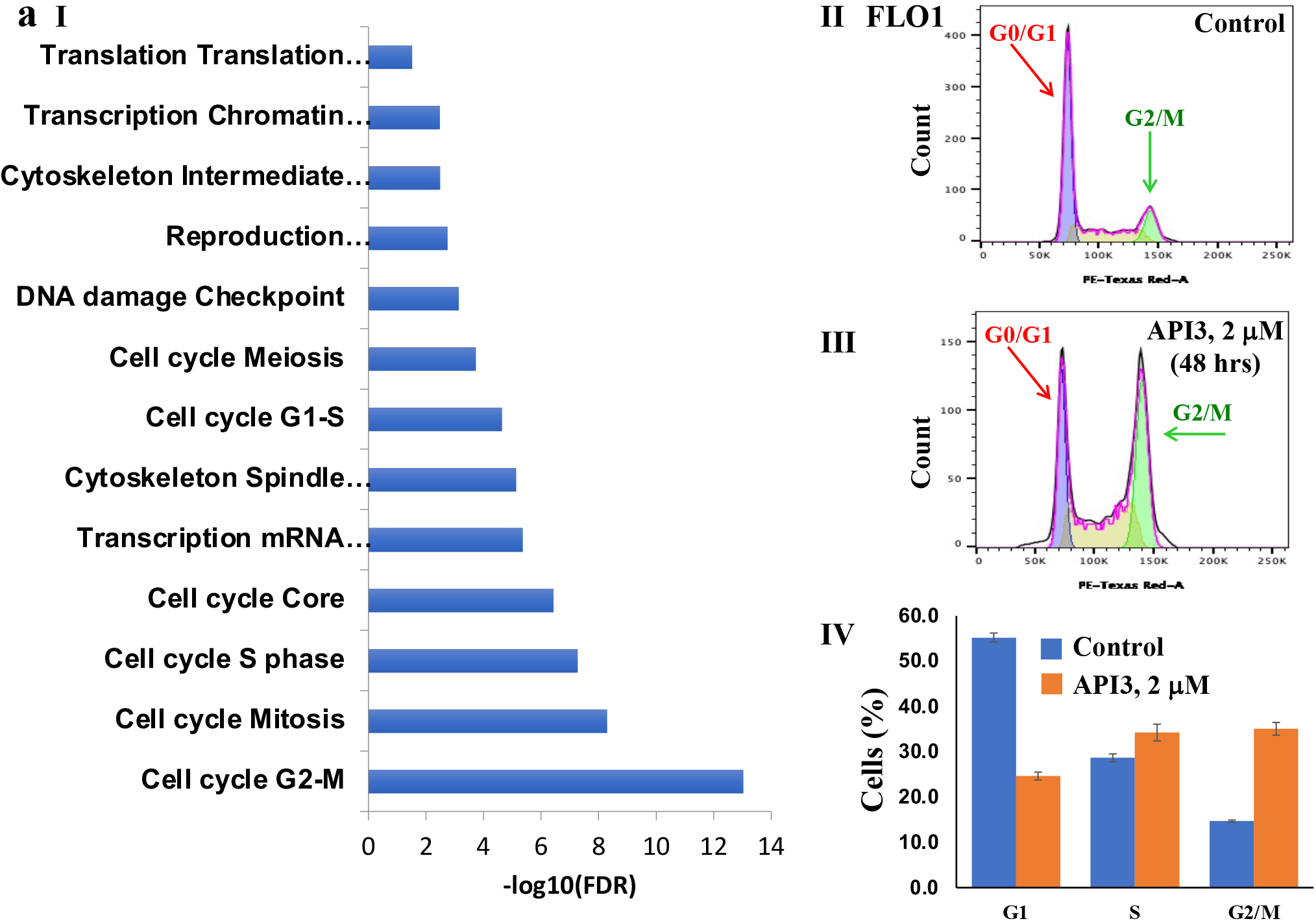
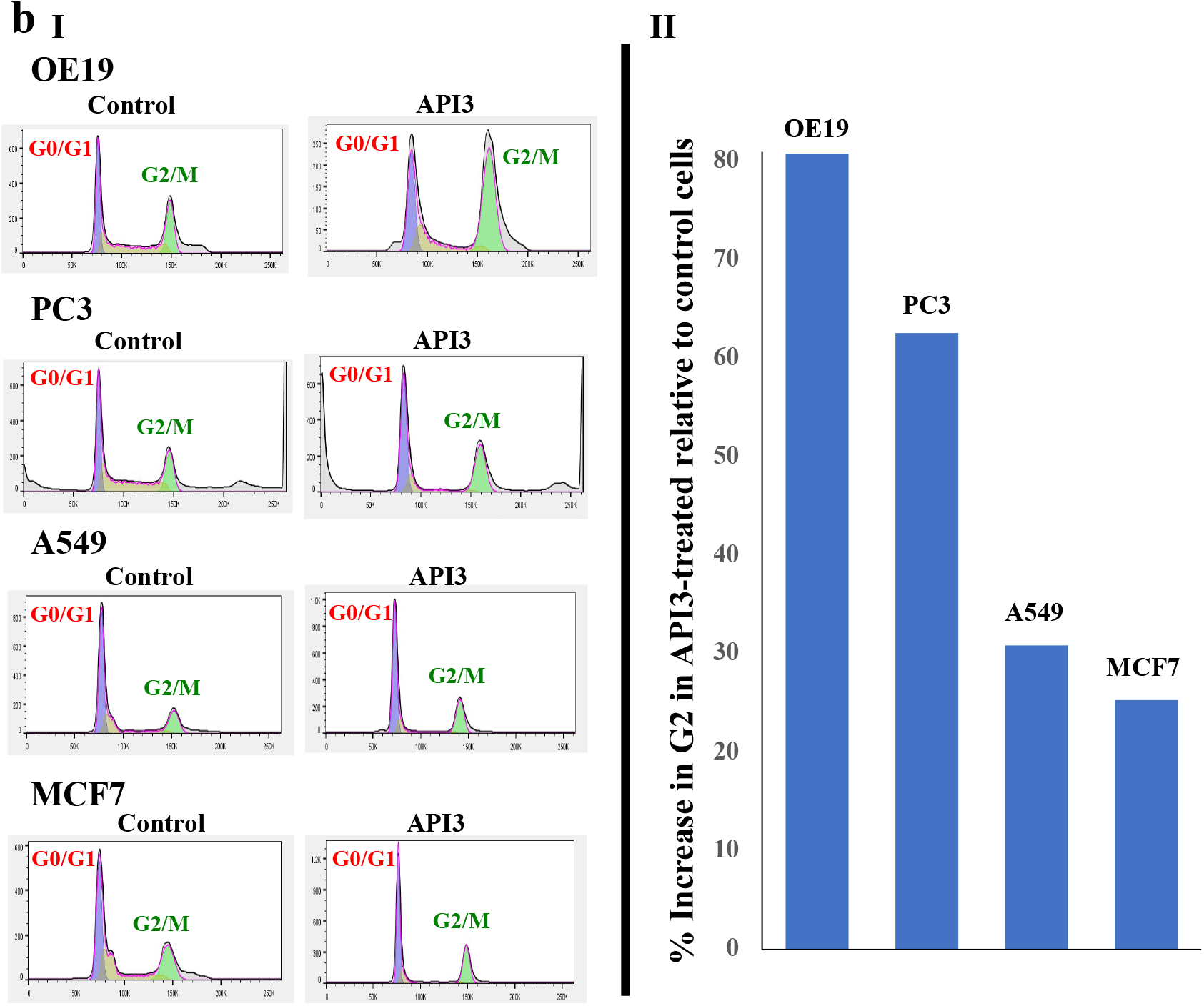
APE1 contributes to G2/M progression in solid cancers. **(A) APE1 regulates G2/M checkpoint in EAC cells. (I)** Expression profile showing common pathways which are upregulated following APE1-overexpression in normal esophageal epithelial (HEsEpi) cells, whereas downregulated in EAC (FLO1) cells treated with APE1 inhibitor (API3); cell cycle (G2/M) is the most significant pathway impacted. **(II-IV)** FLO-1 cells, untreated and treated with API3 (2 ⍰M), were evaluated for cell cycle; cell cycle profiles (II-III) and bar graph showing percentage of cells in different phases of cell cycle (IV) are presented; error bars indicate SDs of triplicate experiment. Two-tailed p value for significance of difference in G2 phase between control and treated cells derived by Student’s t test is 1.5×10^-5^. **(B) Impact of APE1 inhibitor on G2/M checkpoint in different cancers.** Human cancer cell lines - esophageal adenocarcinoma (OE19), breast cancer (MCF7), epithelial lung carcinoma (A549) and prostate adenocarcinoma (PC3), untreated (control) and treated with API3 (1.5 ⍰M), were evaluated for cell cycle; **Panels: (I-IV)** Cell cycle profiles; **(V)** Bar graph showing percent increase in G2 fraction relative to control cells for all cancer cell lines.

## DISCUSSION

Genomic instability, a common feature of precancerous and cancer cells^1–13^, leads to ongoing acquisition of changes at both the DNA and chromosome level. These changes not only provide new characteristics for growth and survival of these cells but also contribute to escape of aberrant cells from immune surveillance^14^, development of resistance to cancer treatment^11,13,15^, progression to advanced stages of disease^63^ and ultimately impact the clinical outcome of disease^13^. A striking genomic instability observed in EAC and its premalignant states^3,8,18–24^ could be attributed to its chemoresistant^25^ nature. A marked genomic instability has also been observed in breast^30^, lung^27^ and prostate^29^ cancers and is attributed to disease progression ^29–30^ and poor clinical outcome^28^. These observations suggest that drivers of genomic instability could serve as promising targets to treat and/or prevent cancer. Consistently, it is now becoming more evident that genomic instability is a promising target in cancer^34,35^. The purpose of this study was to identify genes driving genomic instability in EAC as well as other solid tumors. Since DNA must be cut or broken for genomic rearrangements to take place, we hypothesized that dysregulated deoxyribonuclease activity mediates genomic instability in cancer. Consistent with this hypothesis, the evaluation of **γ**H2AX expression in nine cancer cell lines representing five solid tumors demonstrated that spontaneous DNA breaks are increased in cancer relative to normal cell types (**Supplementary Figure 1**). To find deoxyribonucleases which contribute to spontaneous DNA damage and instability, we used an integrated genomics strategy similar to that described by us previously^54^. Although ribonucleases also contribute to genomic instability, in this study, we focused on deoxyribonucleases because of their ability to directly impact DNA. Since topoisomersases can have additional (helicase and ligase) activities, they were also not included in this analysis. We thus identified a four gene deoxyribonuclease signature correlating with genomic instability in patient (TCGA) datasets representing six solid tumors (adenocarcinomas of esophagus, lung, prostate, stomach, pancreas and triple negative breast cancer). Elevated expression of this signature also correlated with poor overall survival in pancreatic, lung and one of the esophageal adenocarcinoma datasets, establishing the functional relevance of these nucleases. Moreover, functional studies in EAC cells confirmed that inhibition of these nucleases reduced whereas their overexpression increased genomic instability as well as growth rate of these cancer cells. Overall, APE1 was identified as the top gene whose suppression as well as overexpression had the strongest impact on genomic instability and growth of EAC cells. Therefore, APE1 and its inhibitor were investigated in detail for impact on various parameters of genome stability and growth in cancer cell lines representing four different solid cancers including EAC.

APE1 (apurinic/apyrimidinic endonuclease 1), a major base excision repair (BER) enzyme, contributes to the repair process by cleaving the DNA at 5’ of abasic site^45^. Loss of a base or generation of abasic site occurs frequently in a cell^46–48^. Abasic sites in DNA can be produced either spontaneously (by exposure to acid, heat or other factors) in the body or by the activity of DNA glycosylases, the enzymes which remove damaged or altered bases from DNA. DNA glycosylase activity can also be stimulated by APE1 itself. For example, reactive oxygen species can oxidize guanine base leading to formation of 8-oxo-7, 8-dihydroguanine which can incorrectly anneal with adenine causing errors during replication. This aberrant base is removed by a DNA glycosylase called hOGG1 which can be regulated by APE1, the enzyme which cleaves and processes the abasic sites for their repair^64–65^. Abasic sites must be repaired to avoid their mutagenic consequence. In normal cellular environment, thousands of abasic sites are generated daily and efficiently repaired^49,50^. In mammalian cells, APE1 also contributes to 3’-end processing^45^. Moreover, APE1 also has a reduction-oxidation function which is involved in the regulation of transcription factors^66–67^. Thus, APE1 has important roles in the maintenance of genome stability and growth of cells.

Investigation of patient samples in TCGA dataset showed that relative to normal tissues the expression of APE1 is significantly elevated in human cancer specimens of hormone positive and triple negative breast cancers and adenocarcinomas of esophagus (EAC), lung, prostate and colon. Consistently, the evaluation of APE1 expression in frozen tissue specimens of normal squamous epithelium, precancerous Barrett’s esophagus, dysplasia and EAC indicated a markedly elevated expression in dysplasia and EAC relative to that in normal and precancerous cells. Significantly elevated APE1 expression in EAC vs. normal esophageal tissue specimens was also observed in a tissue array. Consistent with our data, elevated APE1 expression has also been reported by other investigators in cervical^68^, ovarian^69^, rhabdomyosarcomas^70^, prostate^71^, and germ cell tumors^72^ thus supporting its potential to be used as a therapeutic target. We, therefore, investigated the impact of APE1 modulation on various parameters of genomic instability, growth, chemoresistance, oncogenic transformation and underlying mechanisms.

Based on our hypothesis that dysregulated nuclease activity increases DNA breaks which lead to genomic instability, we first investigated the impact of APE1 modulation on micronuclei, which is a marker of ongoing DNA/chromosomal rearrangements and instability in a cell^58^. Suppression of APE1, by shRNA-mediated knockdown or treatment of cells with APE1 inhibitor, significantly inhibited genomic instability in all five cancer cell lines representing four solid tumors (esophageal and prostate adenocarcinomas, breast cancer and epithelial lung carcinoma). Evaluation in esophageal adenocarcinoma and epithelial lung carcinoma cell lines demonstrated that treatment with chemotherapeutic agent cisplatin caused further increase in genomic whereas addition of API3 led to near complete inhibition of cisplatin-induced genomic instability in these cell lines. To further evaluate role of APE1 in genomic instability, it was overexpressed in normal esophageal epithelial cells. After sixty days in culture, evaluation by SNP arrays and whole genome sequencing demonstrated that APE1 overexpression in normal cells was associated with a striking increase in the acquisition of copy number and mutational events, respectively. Chromosomes of APE1-overexpressing cells were visualized by spectral karyotyping. The control cells had near diploid karyotype with mitotic index of 0.16% and had 0.2 aberrations/cell, whereas APE1-overexpressing cells were near tetraploid with mitotic index of 3.9%, 16 aberrations/cell, 4 copies of C-myc/cell and several chromosomal abnormalities, indicating a striking karyotypic instability. In summary, using multiple cell and tumor types and evaluation by different platforms including whole genome sequencing, we demonstrate that suppression of APE1 in cancer cells inhibits spontaneous and chemotherapeutic agent-induced genomic instability, whereas its overexpression leads to a striking increase in genomic instability at both DNA and chromosome level. To our knowledge, this is first report demonstrating acquisition of copy number, mutational and karyotypic changes over time by APE1.

Our previous investigation in esophageal adenocarcinoma and multiple myeloma model systems have demonstrated that homologous recombination (HR) is spontaneously elevated and thus dysregulated and attributed to acquisition of genomic rearrangements over time^11–12^, development of resistance to therapy^11^ and growth of cancer cells in vivo^44^. Our investigation in multiple myeloma has also demonstrated that APE1 contributes to increased HR activity through transcriptional regulation of recombinase RAD51^51^. However, dysregulated APE1 activity can also impact HR through induction of DNA breaks. We, therefore, investigated the impact of APE1 modulation on DNA breaks and HR activity in cancer and normal cells. We demonstrate that inhibition of APE1 by its inhibitor or knockdown strongly inhibits cisplatin-induced DNA breaks in EAC and other cancer cell lines. Although APE1 suppression did not seem to impact spontaneous DNA breaks, this could be because under spontaneous condition the number of DNA breaks in these cell lines was quite low or undetectable. However, overexpression of APE1 in normal esophageal epithelial cells led to a substantial increase in spontaneous DNA breaks as assessed by both the Comet assay and Western blotting for **γ**H2AX expression. These data indicate that elevated APE1 increases spontaneous as well as chemotherapeutic agent-induced DNA breaks in affected cells. Inhibition APE1 by its inhibitor or knockdown in EAC cells also led to significant inhibition of RAD51 promoter activity, RAD51 expression and HR activity as assessed by a functional assay. Inhibition APE1 by its inhibitor or knockdown also caused inhibition of HR activity in cell lines representing prostate adenocarcinoma, breast cancer and epithelial lung carcinoma. Moreover, the overexpression of APE1 in normal esophageal cells led to significant increase in HR activity. These data are consistent with our data in multiple myeloma^51^ which show that APE1 contributes to increased HR activity through transcriptional regulation of HR activity. Moreover, identification of APE1-interacting proteins in EAC (FLO-1) cells by mass spectrometry and evaluation of top interactors by protein-protein network analysis identified several DNA repair pathways including those related to HR. These include double-strand break repair, DNA recombination, DNA duplex unwinding, response to gamma radiation, telomere maintenance, base excision repair and non-homologous end joining. Several proteins with known function in HR including RPA1^59^, WRN^60^, DHX9^61^, ILF2^62^ and YBX1^62^ were among top interactors. Furthermore, the evaluation of the impact of APE1-overexpression in normal cells by whole genome sequencing identified HR as the top mutational signature (Figure 5d, III). Since mutational signatures point to the underlying mutational processes^73–74^, these data provide further evidence for the role of HR in APE1-mediated genomic instability. Overall, these data demonstrate that elevated APE1 contributes to increased DNA breaks and HR activity in EAC as well as other solid tumors.

APE1 knockdown led to reduced growth rate in all five cell lines representing four solid tumors (breast cancer, epithelial lung carcinoma and adenocarcinomas of esophagus and prostate). Consistently, the knockdown as well as treatment with APE1 inhibitor, strongly impaired colony formation in all five cell lines. Treatment with APE1 inhibitor also led to a significant increase in cytotoxicity of chemotherapeutic agent cisplatin in all five cell lines. APE1 inhibitor also inhibited tumor growth and increased cytotoxicity of cisplatin in a subcutaneous tumor model of EAC. In line with our observations, a previous report has demonstrated that APE1 contributes to growth of pancreatic cancer cells^75^. Expression of APE1 in cervical cancers has been shown to be commensurate with radiosensitivity^76^. Moreover, a role of APE1 in the regulation of genes involved in chemoresistance has also been proposed^77^. Consistent with APE1 suppression data, its overexpression induced striking genomic instability (at DNA and chromosomal level) and led to increased growth rate and oncogenic transformation of normal esophageal epithelial cells. After around two months in culture, APE1-overexpressing cells grew as tumors in SCID mice whereas control cells did not. Investigation of tumors from euthanized mice by karyotyping identified additional karyotypic changes indicating ongoing genetic instability. These data demonstrate that overexpression of APE1 in normal cells can induce genomic instability predisposing them to oncogenic transformation, whereas its inhibition can impair growth and increase in cytotoxicity of chemotherapeutic agent in EAC and other solid tumors.

Investigation of common pathways impacted by APE1 suppression in EAC and its overexpression in normal esophageal epithelial cells by RNASeq, identified G2/M checkpoint as the most significant common pathway impacted (Figure 7a). Consistently, the treatment with APE1 inhibitor, induced G2/M arrest in all five human cancer cell lines representing four solid tumors. Functional role of APE1 in G2/M progression could be attributed to its ability to impact growth and genomic instability in cancer cells. Cell cycle checkpoints ensure repair of DNA damage prior to DNA replication (in G1) and segregation (in G2), thus ensuring maintenance of genomic integrity. If DNA breaks persist from G_2_ into mitosis, they can recombine in G_1_ to produce gene rearrangements. Moreover, G2 is the phase where HR can utilize sister chromatid as template to ensure precise repair of the damage. Thus, defective G2/M checkpoint^78^ and dysregulated HR^11–12,42^ plays significant role in genomic rearrangements and evolution in cancer.

In this study we demonstrate that inhibition of APE1 reduces spontaneous and cisplatin-induced genomic instability whereas impairs growth and increases cytotoxicity of cisplatin. Here, we would like to emphasize that for all genomic instability related assays, an effort was made to remove dead cells by extensive washing. Our observations can be explained as following: After cisplatin treatment, the cells which acquire extensive DNA damage, are killed. However, the cells which harbor a small or manageable increase in DNA breaks, survive the treatment. Our data indicate that these surviving cells have further increase in HR activity and genomic instability. It seems that dysregulated HR contributes to genomic instability (through aberrant repair of DNA breaks) as well as survival of these cells by reducing DNA damage. Consistently, our data show that inhibition of APE1 reduces chemotherapy-induced genomic instability whereas increases its cytotoxicity. These data are consistent with our previous data in myeloma demonstrating that APE1 inhibitor increases cytotoxicity whereas reduces genomic instability caused by a chemotherapeutic agent^51^. Similar observations have been made following inhibition of HR (by treatment of RAD51 inhibitor) in EAC cells^79^. Our previous data in EAC have also demonstrated that inhibition of genes driving genomic evolution inhibit HR activity, reduce spontaneous and chemotherapy-induced genomic instability, whereas increase cytotoxicity of chemotherapeutic agents^54^.

In summary, we demonstrate the APE1, identified as part of four gene deoxyribonuclease signature, drives genomic instability predisposing normal cells to tumorigenesis. Investigation in five cell lines representing four solid tumors demonstrate that inhibition of APE1 (by knockdown or specific APE1 inhibitor) inhibits spontaneous and chemotherapeutic agent-induced genomic instability, impairs growth/colony formation and increases cytotoxicity of chemotherapeutic agent in cancer cells. Impact of elevated APE1 on DNA breaks and its functional role in dysregulation of G2/M checkpoint and HR could be attributed to genomic instability, growth and chemoresistance in EAC and other solid tumor cell lines tested in this study. Therefore, inhibitors of APE1, alone and/or in combination with other agents, have potential to make EAC and other cancers (including breast, lung and prostate) static at multiple levels.

## References

1 Grady, W. M. Genomic instability and colon cancer. Cancer Metastasis Rev 23, 11–27 (2004).

2 Kawano, M. M. [Genomic instability in multiple myeloma]. Nihon Rinsho 65 Suppl 1, 215–219 (2007).

3 Paulson TG, Maley CC, Li X, Li H, Sanchez CA, Chao DL, Odze RD, Vaughan TL, Blount PL, Reid BJ. Chromosomal instability and copy number alterations in Barrett’s esophagus and esophageal adenocarcinoma. Clin Cancer Res. 2009 May 15;15(10):3305–14. doi: 10.1158/1078-0432.CCR-08-2494. Epub 2009 May 5. PMID: 19417022; PMCID: PMC2684570.

4 Sawyer JR, Tricot G, Lukacs JL, Binz RL, Tian E, Barlogie B, Shaughnessy J Jr. Genomic instability in multiple myeloma: evidence for jumping segmental duplications of chromosome arm 1q. Genes Chromosomes Cancer. 2005 Jan;42(1):95–106. doi: 10.1002/gcc.20109. PMID: 15472896.

5 Sieber, O. M., Heinimann, K. & Tomlinson, I. P. Genomic instability--the engine of tumorigenesis? Nat Rev Cancer 3, 701–708, doi:10.1038/nrc1170 (2003).

6 Timuragaoglu, A., Demircin, S., Dizlek, S., Alanoglu, G. & Kiris, E. Microsatellite instability is a common finding in multiple myeloma. Clin Lymphoma Myeloma 9, 371–374, doi:10.3816/CLM.2009.n.072 (2009).

7. Croft J, Parry EM, Jenkins GJ, Doak SH, Baxter JN, Griffiths AP, Brown TH, Parry JM. Analysis of the premalignant stages of Barrett’s oesophagus through to adenocarcinoma by comparative genomic hybridization. Eur J Gastroenterol Hepatol. 2002 Nov;14(11):1179–86. doi: 10.1097/00042737-200211000-00004. PMID: 12439111.

8 Rabinovitch, P. S., Reid, B. J., Haggitt, R. C., Norwood, T. H. & Rubin, C. E. Progression to cancer in Barrett’s esophagus is associated with genomic instability. Lab Invest 60, 65–71 (1989).

9 Usmani, B. A. Genomic instability and metastatic progression. Pathobiology 61, 109–116, doi:10.1159/000163771 (1993).

10 Bolli N, Avet-Loiseau H, Wedge DC, Van Loo P, Alexandrov LB, Martincorena I, Dawson KJ, Iorio F, Nik-Zainal S, Bignell GR, Hinton JW, Li Y, Tubio JM, McLaren S, O’ Meara S, Butler AP, Teague JW, Mudie L, Anderson E, Rashid N, Tai YT, Shammas MA, Sperling AS, Fulciniti M, Richardson PG, Parmigiani G, Magrangeas F, Minvielle S, Moreau P, Attal M, Facon T, Futreal PA, Anderson KC, Campbell PJ, Munshi NC. Heterogeneity of genomic evolution and mutational profiles in multiple myeloma. Nat Commun. 2014;5:2997. doi: 10.1038/ncomms3997. PMID: 24429703; PMCID: PMC3905727.

11 Shammas MA, Shmookler Reis RJ, Koley H, Batchu RB, Li C, Munshi NC. Dysfunctional homologous recombination mediates genomic instability and progression in myeloma. Blood. 2009 Mar 5;113(10):2290–7. doi: 10.1182/blood-2007-05-089193. Epub 2008 Dec 2. PMID: 19050310; PMCID: PMC2652372.

12 Pal J, Bertheau R, Buon L, Qazi A, Batchu RB, Bandyopadhyay S, Ali-Fehmi R, Beer DG, Weaver DW, Shmookler Reis RJ, Goyal RK, Huang Q, Munshi NC, Shammas MA. Genomic evolution in Barrett’s adenocarcinoma cells: critical roles of elevated hsRAD51, homologous recombination and Alu sequences in the genome. Oncogene. 2011 Aug 18;30(33):3585–98. doi: 10.1038/onc.2011.83. Epub 2011 Mar 21. PMID: 21423218; PMCID: PMC3406293.

13 Bakhoum SF, Cantley LC. The Multifaceted Role of Chromosomal Instability in Cancer and Its Microenvironment. Cell. 2018 Sep 6;174(6):1347–1360. doi: 10.1016/j.cell.2018.08.027. PMID: 30193109; PMCID: PMC6136429.

14 Nandi B, Talluri S, Kumar S, Yenumula C, Gold JS, Prabhala R, Munshi NC, Shammas MA. The roles of homologous recombination and the immune system in the genomic evolution of cancer. J Transl Sci. 2019 Apr;5(2):10.15761/JTS.1000282. doi: 10.15761/JTS.1000282. Epub 2018 Oct 1. PMID: 30873294; PMCID: PMC6411307.

15 Bakhoum SF, Ngo B, Laughney AM, Cavallo JA, Murphy CJ, Ly P, Shah P, Sriram RK, Watkins TBK, Taunk NK, Duran M, Pauli C, Shaw C, Chadalavada K, Rajasekhar VK, Genovese G, Venkatesan S, Birkbak NJ, McGranahan N, Lundquist M, LaPlant Q, Healey JH, Elemento O, Chung CH, Lee NY, Imielenski M, Nanjangud G, Pe’er D, Cleveland DW, Powell SN, Lammerding J, Swanton C, Cantley LC. Chromosomal instability drives metastasis through a cytosolic DNA response. Nature. 2018 Jan 25;553(7689):467–472. doi: 10.1038/nature25432. Epub 2018 Jan 17. PMID: 29342134; PMCID: PMC5785464.

16 Dagogo-Jack I, Shaw AT. Tumour heterogeneity and resistance to cancer therapies. Nat Rev Clin Oncol. 2018 Feb;15(2):81–94.

17 Marusyk A, Janiszewska M, Polyak K. Intratumor Heterogeneity: The Rosetta Stone of Therapy Resistance. Cancer Cell. 2020 Apr 13;37(4):471–484.

18 Spechler SJ, Goyal RK. Barrett’s esophagus. N Engl J Med. 1986 Aug 7;315(6):362–71. doi: 10.1056/NEJM198608073150605. PMID: 2874485.

19 Ross-Innes CS, Becq J, Warren A, Cheetham RK, Northen H, O’Donovan M, Malhotra S, di Pietro M, Ivakhno S, He M, Weaver JMJ, Lynch AG, Kingsbury Z, Ross M, Humphray S, Bentley D, Fitzgerald RC. Whole-genome sequencing provides new insights into the clonal architecture of Barrett’s esophagus and esophageal adenocarcinoma. Nat Genet. 2015 Sep;47(9):1038–1046. doi: 10.1038/ng.3357. Epub 2015 Jul 20. PMID: 26192915; PMCID: PMC4556068.

20 Stachler MD, Taylor-Weiner A, Peng S, McKenna A, Agoston AT, Odze RD, Davison JM, Nason KS, Loda M, Leshchiner I, Stewart C, Stojanov P, Seepo S, Lawrence MS, Ferrer-Torres D, Lin J, Chang AC, Gabriel SB, Lander ES, Beer DG, Getz G, Carter SL, Bass AJ. Paired exome analysis of Barrett’s esophagus and adenocarcinoma. Nat Genet. 2015 Sep;47(9):1047–55. doi: 10.1038/ng.3343. Epub 2015 Jul 20. PMID: 26192918; PMCID: PMC4552571.

21 Gregson EM, Bornschein J, Fitzgerald RC. Genetic progression of Barrett’s oesophagus to oesophageal adenocarcinoma. Br J Cancer 2016;115:403–10.

22 Akagi T, Ito T, Kato M, Jin Z, Cheng Y, Kan T, Yamamoto G, Olaru A, Kawamata N, Boult J, Soukiasian HJ, Miller CW, Ogawa S, Meltzer SJ, Koeffler HP. Chromosomal abnormalities and novel disease-related regions in progression from Barrett’s esophagus to esophageal adenocarcinoma. Int J Cancer. 2009 Nov 15;125(10):2349–59. doi: 10.1002/ijc.24620. PMID: 19670330; PMCID: PMC2766567.

23 Yu C, Zhang X, Huang Q, Klein M, Goyal RK. High-fidelity DNA histograms in neoplastic progression in Barrett’s esophagus. Lab Invest. 2007 May;87(5):466–72. doi: 10.1038/labinvest.3700531. Epub 2007 Feb 19. PMID: 17310216.

24 Cai JC, Liu D, Liu KH, Zhang HP, Zhong S, Xia NS. Microsatellite alterations in phenotypically normal esophageal squamous epithelium and metaplasia-dysplasia-adenocarcinoma sequence. World J Gastroenterol. 2008 Jul 7;14(25):4070–6. doi: 10.3748/wjg.14.4070. PMID: 18609693; PMCID: PMC2725348.

25 Testa U, Castelli G, Pelosi E. Esophageal Cancer: Genomic and Molecular Characterization, Stem Cell Compartment and Clonal Evolution. Medicines (Basel). 2017 Sep 14;4(3):67. doi: 10.3390/medicines4030067. PMID: 28930282; PMCID: PMC5622402.

26 Maret-Ouda J, El-Serag HB, Lagergren J. Opportunities for Preventing Esophageal Adenocarcinoma. Cancer Prev Res (Phila) 2016;9:828–834.

27 Soca-Chafre G, Montiel-Dávalos A, Rosa-Velázquez IA, Caro-Sánchez CHS, Peña-Nieves A, Arrieta O. Multiple Molecular Targets Associated with Genomic Instability in Lung Cancer. Int J Genomics. 2019 Jul 1;2019:9584504. doi: 10.1155/2019/9584504. PMID: 31355244; PMCID: PMC6636528.

28 Geng W, Lv Z, Fan J, Xu J, Mao K, Yin Z, Qing W, Jin Y. Identification of the Prognostic Significance of Somatic Mutation-Derived LncRNA Signatures of Genomic Instability in Lung Adenocarcinoma. Front Cell Dev Biol. 2021 Mar 29;9:657667. doi: 10.3389/fcell.2021.657667. PMID: 33855028; PMCID: PMC8039462.

29 Tapia-Laliena MA, Korzeniewski N, Hohenfellner M, Duensing S. High-risk prostate cancer: a disease of genomic instability. Urol Oncol. 2014 Nov;32(8):1101–7. doi: 10.1016/j.urolonc.2014.02.005. Epub 2014 Jun 13. PMID: 24931269.

30 Kalimutho M, Nones K, Srihari S, Duijf PHG, Waddell N, Khanna KK. Patterns of Genomic Instability in Breast Cancer. Trends Pharmacol Sci. 2019 Mar;40(3):198–211. doi: 10.1016/j.tips.2019.01.005. Epub 2019 Feb 6. PMID: 30736983.

31 Duijf PHG, Nanayakkara D, Nones K, Srihari S, Kalimutho M, Khanna KK. Mechanisms of Genomic Instability in Breast Cancer. Trends Mol Med. 2019 Jul;25(7):595–611. doi: 10.1016/j.molmed.2019.04.004. Epub 2019 May 8. PMID: 31078431.

32 Thakkar JP, Mehta DG. A review of an unfavorable subset of breast cancer: estrogen receptor positive progesterone receptor negative. Oncologist. 2011;16(3):276–85. doi: 10.1634/theoncologist.2010-0302. Epub 2011 Feb 21. PMID: 21339261; PMCID: PMC3228102.

33 King L, Flaus A, Holian E, Golden A. Survival outcomes are associated with genomic instability in luminal breast cancers. PLoS One. 2021 Feb 3;16(2):e0245042. doi: 10.1371/journal.pone.0245042. PMID: 33534788; PMCID: PMC7857737.

34 Salmaninejad A, Ilkhani K, Marzban H, Navashenaq JG, Rahimirad S, Radnia F, Yousefi M, Bahmanpour Z, Azhdari S, Sahebkar A. Genomic Instability in Cancer: Molecular Mechanisms and Therapeutic Potentials. Curr Pharm Des. 2021;27(28):3161–3169. doi: 10.2174/1381612827666210426100206. PMID: 33902409.

35 Bielski, C.M., Taylor, B.S. Homing in on genomic instability as a therapeutic target in cancer. Nat Commun 12, 3663 (2021).

36 Solomon E, Borrow J, Goddard AD. Chromosome aberrations and cancer. Science. 1991;254:1153–1160. doi: 10.1126/science.1957167.

37 Tubbs A, Nussenzweig A. Endogenous DNA Damage as a Source of Genomic Instability in Cancer. Cell. 2017 Feb 9;168(4):644–656. doi: 10.1016/j.cell.2017.01.002. PMID: 28187286; PMCID: PMC6591730.

38 Renkawitz J, Lademann CA, Jentsch S. Mechanisms and principles of homology search during recombination. Nat Rev Mol Cell Biol. 2014 Jun;15(6):369–83. doi: 10.1038/nrm3805. Epub 2014 May 14. PMID: 24824069.

39 Heyer WD, Ehmsen KT, Liu J. Regulation of homologous recombination in eukaryotes. Annu. Rev. Genet. 2010;44:113–139. doi: 10.1146/annurev-genet-051710-150955.

40 Moynahan ME, Jasin M. Mitotic homologous recombination maintains genomic stability and suppresses tumorigenesis. Nat. Rev. Mol. Cell Biol. 2010;11:196–207. doi: 10.1038/nrm2851.

41 Orthwein A, Noordermeer SM, Wilson MD, Landry S, Enchev RI, et al. (2015) A mechanism for the suppression of homologous recombination in G1 cells. Nature 528: 422–426.

42 Guirouilh-Barbat J, Lambert S, Bertrand P, Lopez BS (2014) Is homologous recombination really an error-free process? Front Genet 5: 170–175.

43 Branzei D, Foiani M (2008) 1Regulation of DNA repair throughout the cell cycle. Nat Rev Mol Cell Biol 9:297–308.

44 Lu, R. et al. Targeting homologous recombination and telomerase in Barrett’s adenocarcinoma: impact on telomere maintenance, genomic instability and tumor growth. Oncogene 33, 1495–1505, doi:10.1038/onc.2013.103 (2014).

45 Mol CD, Hosfield DJ, Tainer JA. Abasic site recognition by two apurinic/apyrimidinic endonuclease families in DNA base excision repair: the 3’ ends justify the means. Mutat. Res. 2000;460:211–229. doi: 10.1016/S0921-8777(00)00028-8.

46 Barzilay G, Hickson ID. Structure and function of apurinic/apyrimidinic endonucleases. Bioessays. 1995;17:713–719. doi: 10.1002/bies.950170808.

47 Barzilay G, Walker LJ, Rothwell DG, Hickson ID. Role of the HAP1 protein in repair of oxidative DNA damage and regulation of transcription factors. Br. J. Cancer Suppl. 1996;27:S145–S150.

48 Rothwell DG, et al. The structure and functions of the HAP1/Ref-1 protein. Oncol. Res. 1997;9:275–280.

49 Frosina G, Fortini P, Rossi O, Carrozzino F, Abbondandolo A, Dogliotti E. Repair of abasic sites by mammalian cell extracts. Biochem J. 1994 Dec 15;304 (Pt 3)(Pt 3):699–705. doi: 10.1042/bj3040699. PMID: 7818470; PMCID: PMC1137391.

50 Nakamura J, Swenberg JA. Endogenous apurinic/apyrimidinic sites in genomic DNA of mammalian tissues. Cancer Res. 1999 Jun 1;59(11):2522–6. PMID: 10363965.

51 Kumar S, Talluri S, Pal J, Yuan X, Lu R, Nanjappa P, Samur MK, Munshi NC, Shammas MA. Role of apurinic/apyrimidinic nucleases in the regulation of homologous recombination in myeloma: mechanisms and translational significance. Blood Cancer J. 2018 Sep 25;8(10):92. doi: 10.1038/s41408-018-0129-9. PMID: 30301882; PMCID: PMC6177467.

52 Pal J, Nanjappa P, Kumar S, et al. Impact of RAD51C-mediated Homologous Recombination on Genomic Integrity in Barrett’s Adenocarcinoma Cells. J Gastroenterol Hepatol Res 2017; 6:2286–2295.

53 Shammas MA, Koley H, Beer DG, et al. Growth arrest, apoptosis, and telomere shortening of Barrett’s-associated adenocarcinoma cells by a telomerase inhibitor. Gastroenterology 2004; 126:1337–46.

54 Kumar S, Buon L, Talluri S, Roncador M, Liao C, Zhao J, Shi J, Chakraborty C, Gonzalez G, Tai YT, Prabhala R, Samur MK, Munshi NC, Shammas MA. Integrated genomics and comprehensive validation reveal drivers of genomic evolution in esophageal adenocarcinoma. Commun Biol. 2021 May 24;4(1):617. doi: 10.1038/s42003-021-02125-x. PMID: 34031527; PMCID: PMC8144613.

55 Huang F, Motlekar NA, Burgwin CM, Napper AD, Diamond SL, Mazin AV. Identification of specific inhibitors of human RAD51 recombinase using high-throughput screening. ACS Chem Biol. 2011 Jun 17;6(6):628–35. doi: 10.1021/cb100428c. Epub 2011 Mar 23. PMID: 21428443; PMCID: PMC3117970.

56 Gravel S, Chapman JR, Magill C, Jackson SP. DNA helicases Sgs1 and BLM promote DNA doublestrand break resection. Genes Dev. 2008;22(20):2767–2772.

57 Sartori AA, Lukas C, Coates J, et al. Human CtIP promotes DNA end resection. Nature. 2007;450(7169):509–514.

58 Balmus G, Karp NA, Ng BL, Jackson SP, Adams DJ, McIntyre RE. A high-throughput in vivo micronucleus assay for genome instability screening in mice. Nat Protoc. 2015 Jan;10(1):205–15. doi: 10.1038/nprot.2015.010. Epub 2014 Dec 31. PMID: 25551665; PMCID: PMC4806852.

59 Toh M, Ngeow J. Homologous Recombination Deficiency: Cancer Predispositions and Treatment Implications. Oncologist. 2021 Sep;26(9):e1526–e1537. doi: 10.1002/onco.13829. Epub 2021 Jun 2. PMID: 34021944; PMCID: PMC8417864.

60 Saintigny Y, Makienko K, Swanson C, Emond MJ, Monnat RJ Jr. Homologous recombination resolution defect in werner syndrome. Mol Cell Biol. 2002 Oct;22(20):6971–8. doi: 10.1128/MCB.22.20.6971-6978.2002. PMID: 12242278; PMCID: PMC139822.

61 Chakraborty P, Hiom K. DHX9-dependent recruitment of BRCA1 to RNA promotes DNA end resection in homologous recombination. Nat Commun. 2021 Jul 5;12(1):4126. doi: 10.1038/s41467-021-24341-z. PMID: 34226554; PMCID: PMC8257769.

62 Marchesini M, Ogoti Y, Fiorini E, Aktas Samur A, Nezi L, D’Anca M, Storti P, Samur MK, Ganan-Gomez I, Fulciniti MT, Mistry N, Jiang S, Bao N, Marchica V, Neri A, Bueso-Ramos C, Wu CJ, Zhang L, Liang H, Peng X, Giuliani N, Draetta G, Clise-Dwyer K, Kantarjian H, Munshi N, Orlowski R, Garcia-Manero G, DePinho RA, Colla S. ILF2 Is a Regulator of RNA Splicing and DNA Damage Response in 1q21-Amplified Multiple Myeloma. Cancer Cell. 2017 Jul 10;32(1):88–100.e6. doi: 10.1016/j.ccell.2017.05.011. Epub 2017 Jun 29. PMID: 28669490; PMCID: PMC5593798.

63 McClelland SE. Role of chromosomal instability in cancer progression. Endocr Relat Cancer. 2017 Sep;24(9):T23–T31. doi: 10.1530/ERC-17-0187. Epub 2017 Jul 10. PMID: 28696210.

64 Vidal AE, Hickson ID, Boiteux S, Radicella JP. Mechanism of stimulation of the DNA glycosylase activity of hOGG1 by the major human AP endonuclease: bypass of the AP lyase activity step. Nucleic Acids Res. 2001 Mar 15;29(6):1285–92. doi: 10.1093/nar/29.6.1285. PMID: 11238994; PMCID: PMC29755.

65 Hill JW, Hazra TK, Izumi T, Mitra S. Stimulation of human 8-oxoguanine-DNA glycosylase by AP-endonuclease: potential coordination of the initial steps in base excision repair. Nucleic Acids Res. 2001 Jan 15;29(2):430–8. doi: 10.1093/nar/29.2.430. PMID: 11139613; PMCID: PMC29662.

66 Fung H, Demple B. A vital role for ape1/ref1 protein in repairing spontaneous DNA damage in human cells. Mol. Cell. 2005;17:463–470. doi: 10.1016/j.molcel.2004.12.029.

67 Kelley MR, Parsons SH. Redox regulation of the DNA repair function of the human AP endonuclease Ape1/ref-1. Antioxid. Redox Signal. 2001;3:671–683. doi: 10.1089/15230860152543014.

68 Schindl M, Oberhuber G, Pichlbauer EG, Obermair A, Birner P, Kelley MR. DNA repair-redox enzyme apurinic endonuclease in cervical cancer: evaluation of redox control of HIF-1alpha and prognostic significance. Int J Oncol. 2001 Oct;19(4):799–802. doi: 10.3892/ijo.19.4.799. PMID: 11562758.

69 Tanner B, Grimme S, Schiffer I, Heimerdinger C, Schmidt M, Dutkowski P, Neubert S, Oesch F, Franzen A, Kölbl H, Fritz G, Kaina B, Hengstler JG. Nuclear expression of apurinic/apyrimidinic endonuclease increases with progression of ovarian carcinomas. Gynecol Oncol. 2004 Feb;92(2):568–77. doi: 10.1016/j.ygyno.2003.10.037. PMID: 14766249.

70 Thomson B, Tritt R, Davis M, Kelley MR. Histology-specific expression of a DNA repair protein in pediatric rhabdomyosarcomas. J Pediatr Hematol Oncol. 2001 May;23(4):234–9. doi: 10.1097/00043426-200105000-00011. Erratum in: J Pediatr Hematol Oncol 2001 Aug-Sep;23(6):369. PMID: 11846302.

71 Kelley MR, Cheng L, Foster R, Tritt R, Jiang J, Broshears J, Koch M. Elevated and altered expression of the multifunctional DNA base excision repair and redox enzyme Ape1/ref-1 in prostate cancer. Clin Cancer Res. 2001 Apr;7(4):824–30. PMID: 11309329.

72 Robertson KA, Bullock HA, Xu Y, Tritt R, Zimmerman E, Ulbright TM, Foster RS, Einhorn LH, Kelley MR. Altered expression of Ape1/ref-1 in germ cell tumors and overexpression in NT2 cells confers resistance to bleomycin and radiation. Cancer Res. 2001 Mar 1;61(5):2220–5. PMID: 11280790.

73 Alexandrov, L. B. et al. Signatures of mutational processes in human cancer. Nature 500, 415–421, doi:10.1038/nature12477 (2013).

74 Alexandrov, L. B., Nik-Zainal, S., Wedge, D. C., Campbell, P. J. & Stratton, M. R. Deciphering signatures of mutational processes operative in human cancer. Cell Rep 3, 246–259, doi:10.1016/j.celrep.2012.12.008 (2013).

75 Choi YD, Jung JY, Baek M, Khan S, Song PI, Ryu S, Koo JY, Chauhan SC, Tsin A, Choi C, Kim WJ, Kim M. APE1 Promotes Pancreatic Cancer Proliferation through GFRα1/Src/ERK Axis-Cascade Signaling in Response to GDNF. Int J Mol Sci. 2020 May 19;21(10):3586. doi: 10.3390/ijms21103586. PMID: 32438692; PMCID: PMC7279477.

76 Herring, C., West, C., Wilks, D. et al. Levels of the DNA repair enzyme human apurinic/apyrimidinic endonuclease (APE1, APEX, Ref-1) are associated with the intrinsic radiosensitivity of cervical cancers. Br J Cancer 78, 1128–1133 (1998). https://doi.org/10.1038/bjc.1998.641

77 Ayyildiz, D., Antoniali, G., D’Ambrosio, C. et al. Architecture of The Human Ape1 Interactome Defines Novel Cancers Signatures. Sci Rep 10, 28 (2020).

78 Kops, G., Weaver, B. & Cleveland, D. On the road to cancer: aneuploidy and the mitotic checkpoint. Nat Rev Cancer 5, 773–785 (2005).

79 Liao C, Zhao J, Kumar S, Chakraborty C, Talluri S, Munshi NC, Shammas MA. RAD51 Inhibitor Reverses Etoposide-Induced Genomic Toxicity and Instability in Esophageal Adenocarcinoma Cells. Arch Clin Toxicol (Middlet). 2020;2(1):3–9. doi: 10.46439/toxicology.2.006. PMID: 32968740; PMCID: PMC7508453.

