## Supplementary Figures for "Apurinic/apyrimidinic nuclease 1 drives genomic evolution contributing to chemoresistance and tumorigenesis in solid tumor"

**Supplementary Figure 1**

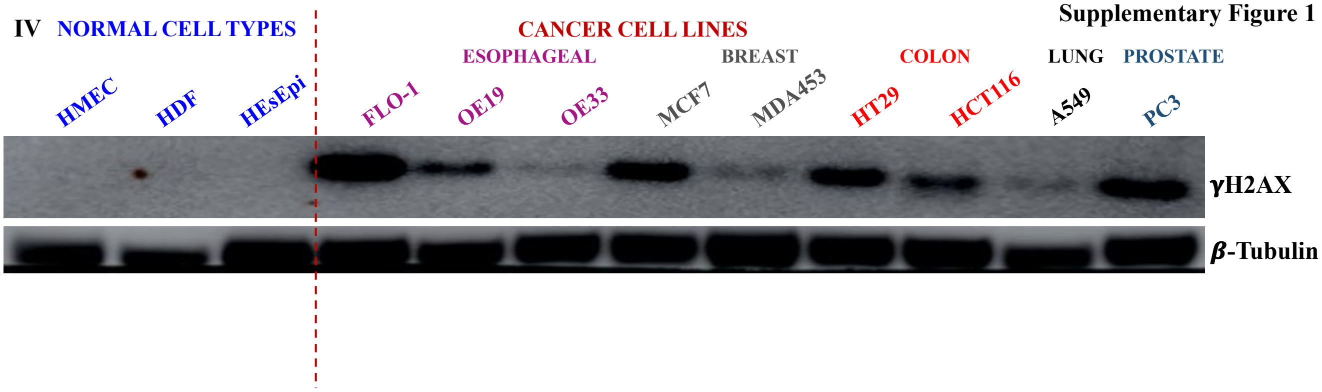


**Supplementary Figure 2**

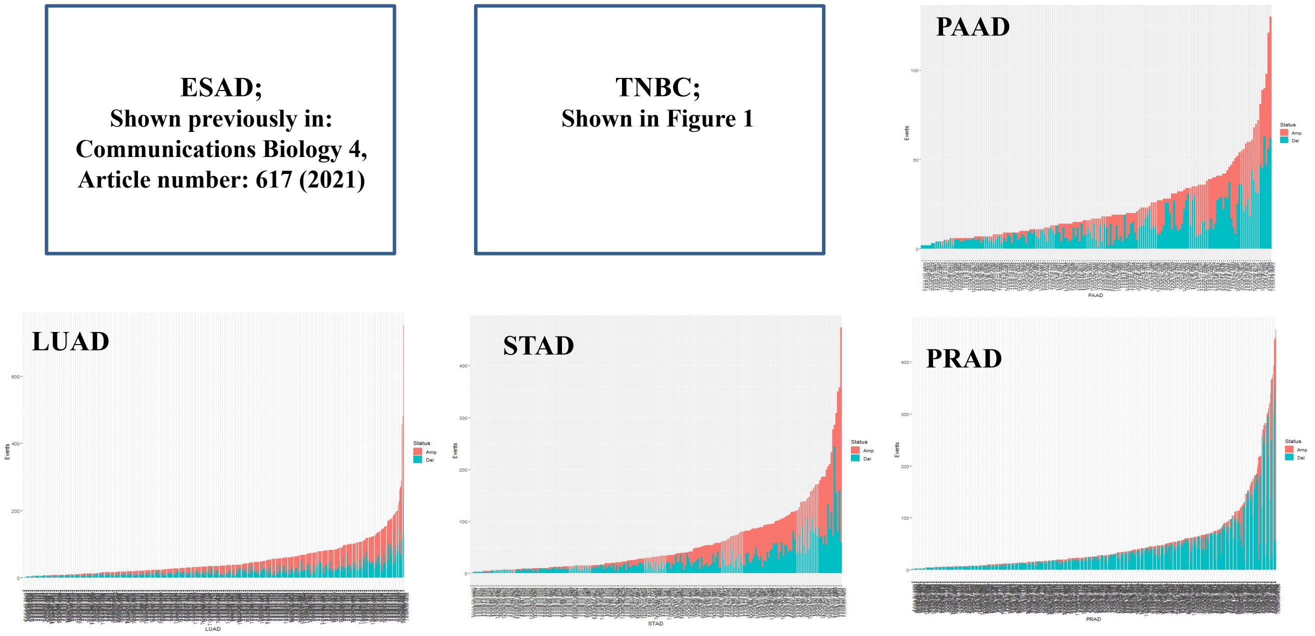

Supplementary Figure 3

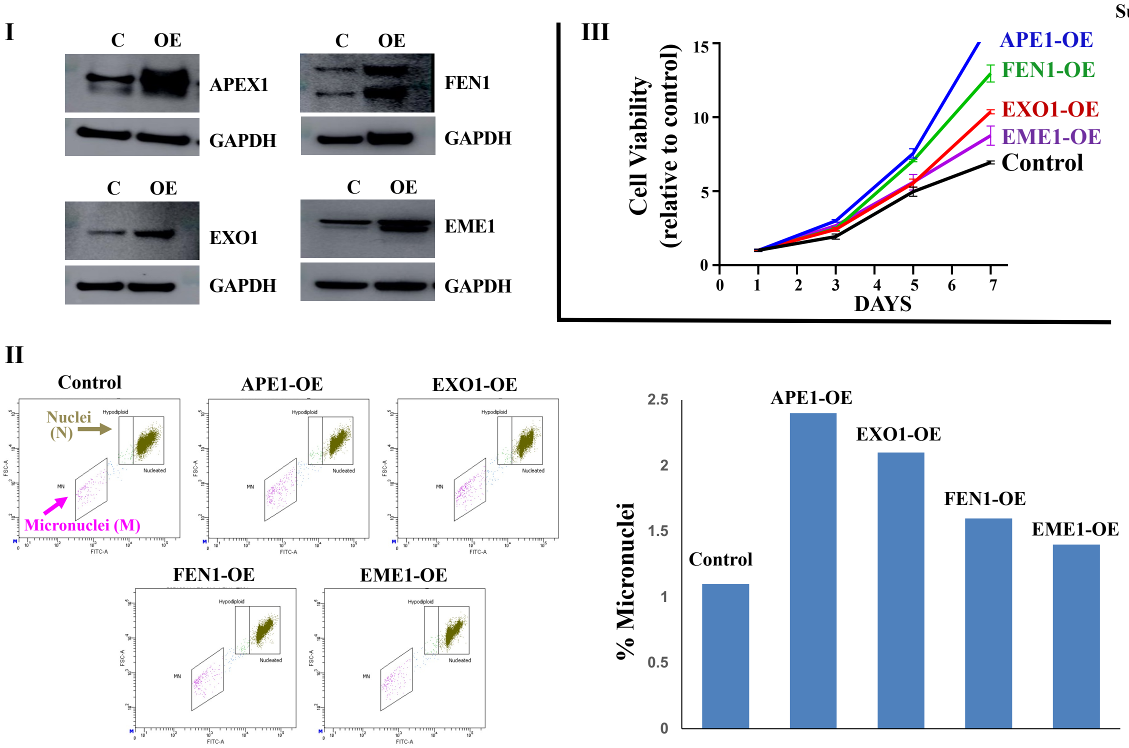


Supplementary Figure 4

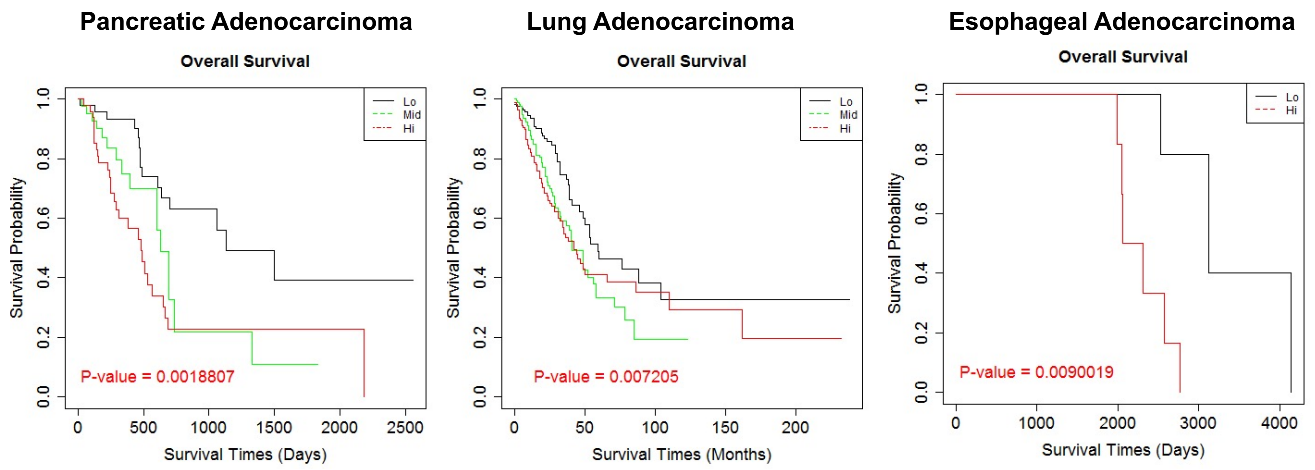

**Supplementary Figure 5**

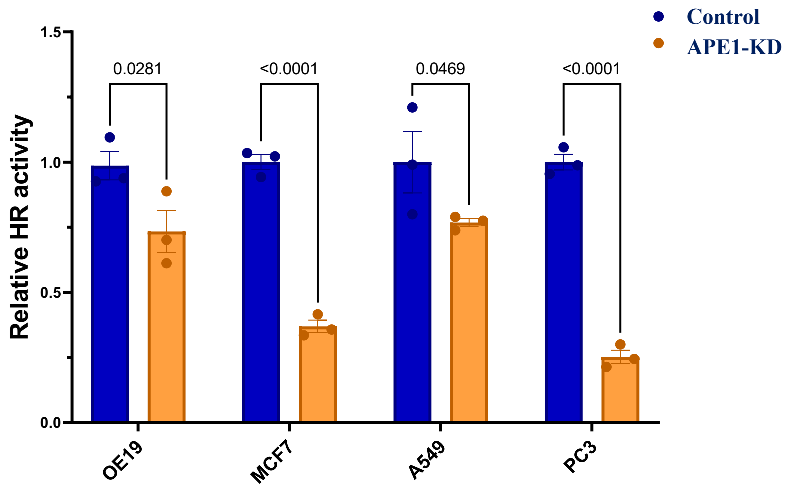


**Supplementary Figure 6**

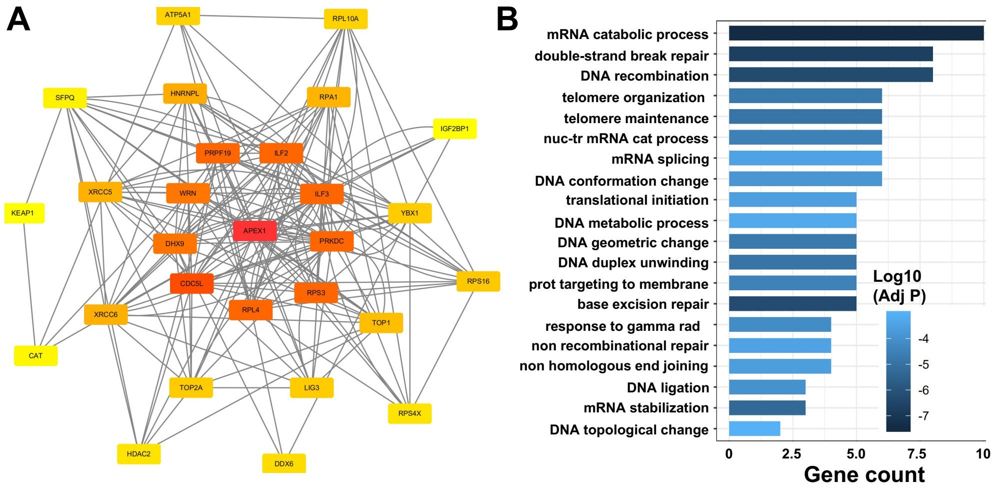

**a** **Supplementary Figure 7**

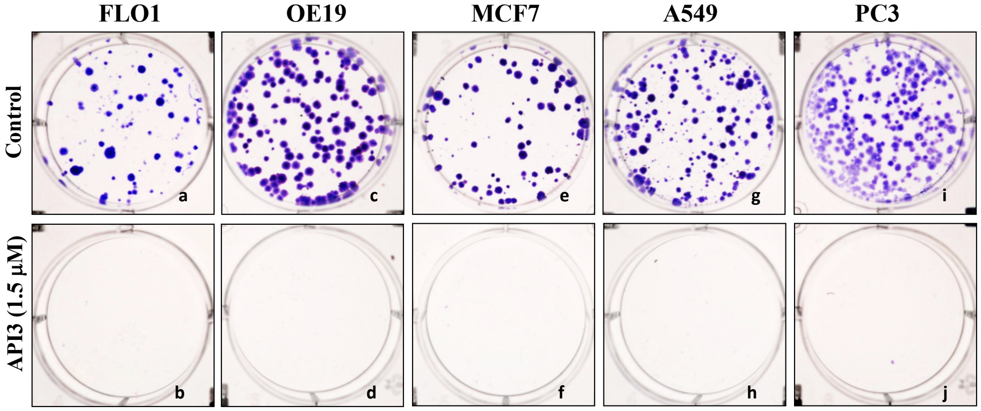


**Supplementary Figure 7**

**b**

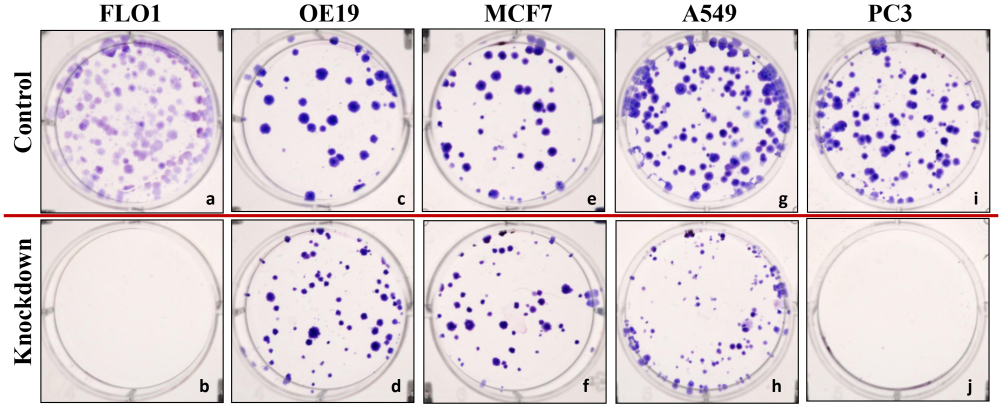

Supplementary Figure 8

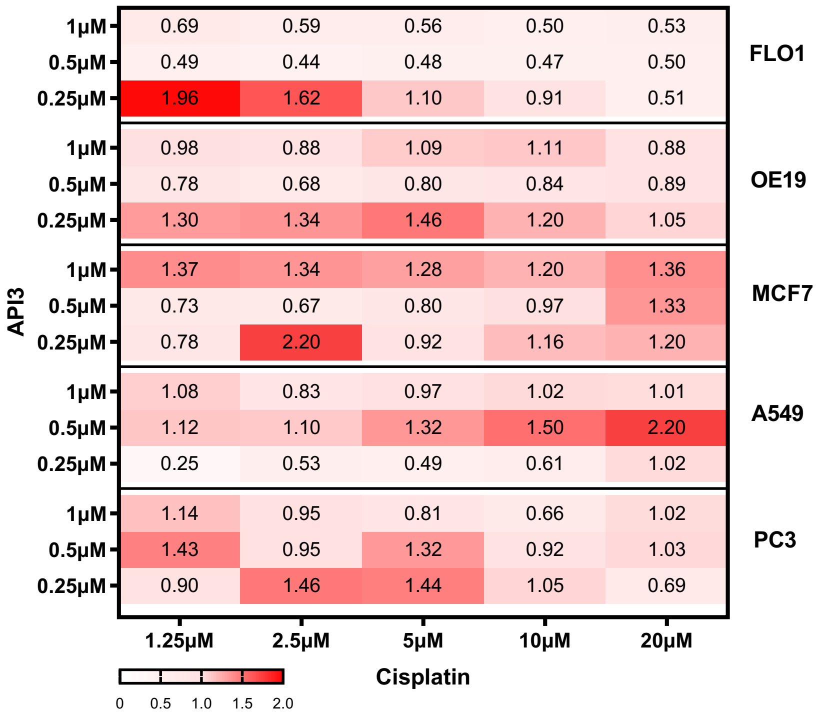


CI<1; Synergism
CI=1: Additive Effect
CI >1: Antagonism

Supplementary Figure 9

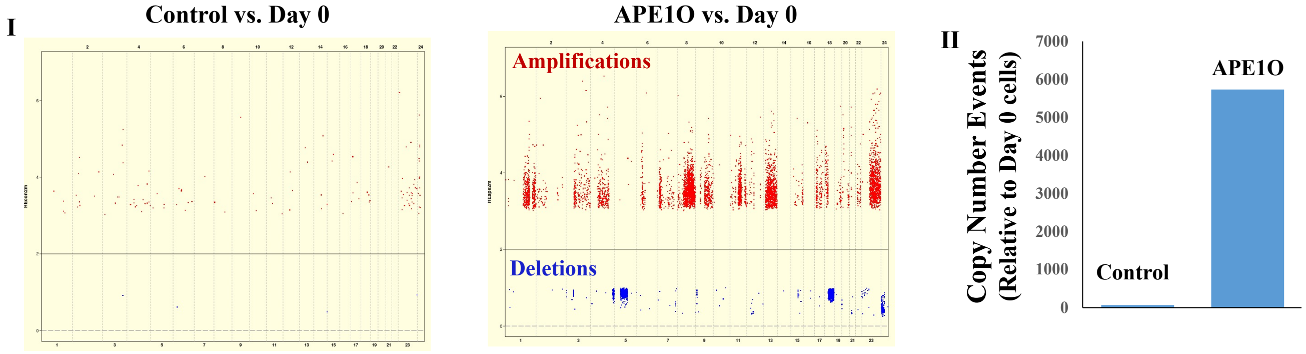

### Supplementary Figure 10

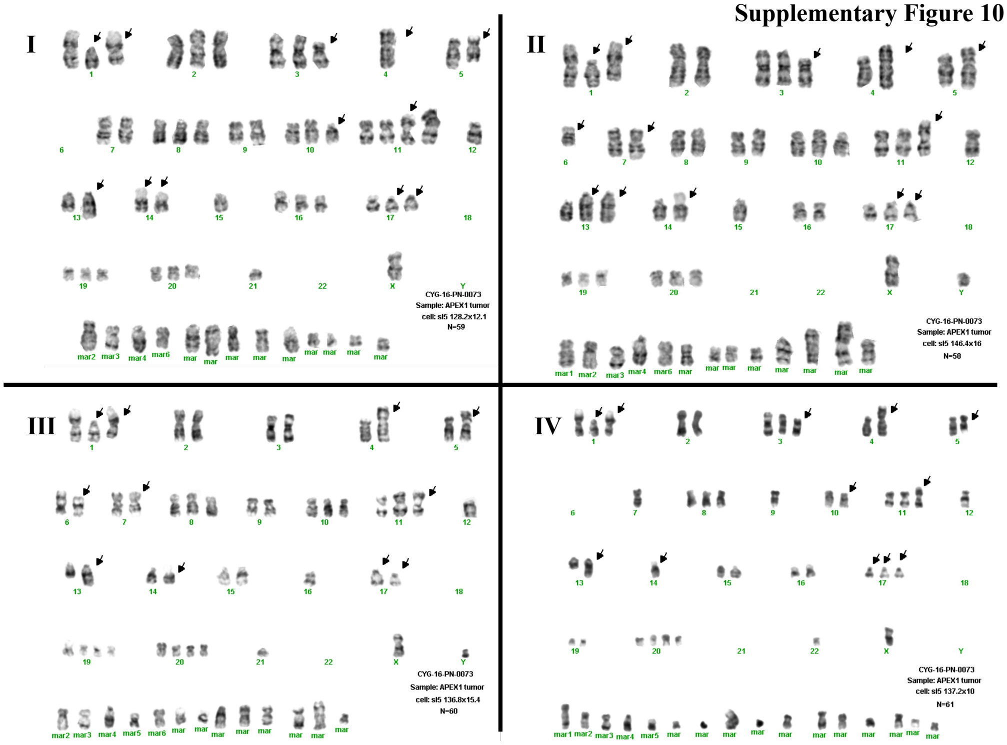
